# DigitAb: Domain-Adaptive Cell Type Prediction Method from Light Microscopy Images

**DOI:** 10.64898/2026.05.19.726313

**Authors:** Nicholas Lucarelli, Seth Winfree, Angela Sabo, Daria Barwinska, Michael Ferkowicz, William Bowen, Akshat Singh, Kaifeng Chen, Anish Tatke, Kuang-Yu Jen, Michael T Eadon, Tarek M. El-Achkar, Sanjay Jain, Pinaki Sarder

## Abstract

Light microscopy imaging with histological stains is central to disease diagnosis and research. It is enhanced with immunostaining to reveal cellular composition and complexity linked to clinical utility and biological mechanisms. Emerging multiplex imaging technologies like Phenocycler markedly increase the coverage to capture the cellular diversity but are costly, technically demanding, and inaccessible to most clinical laboratories. We developed DigitAb, a deep learning framework that classifies cell types directly from hematoxylin and eosin (H&E) stained slides, eliminating the need for specialized assays. Using Phenocycler imaging, we generated highlZlresolution ground truths for ∼3.5 million cells from 29 human kidney samples across four multi-institutional datasets to train a semantic segmentation model for 10 cell types, achieving a balanced accuracy of 0.78. By employing an integrated adversarial domain adaptation module, we tested DigitAb on unlabeled and untested biopsy samples from kidney transplant and diabetic samples. We were able to predict several cell types just from histology images, without using any special technology or immunostains, and demonstrate high concordance with clinical gold-standard Banff schema in kidney transplant rejection, and clinical characteristics of diabetic nephropathy. Our cloudlZlbased tool, DigitAb, provides scalable, accessible, labellZlfree cellular segmentation for research and clinical pathology.

## Main

Studying tissue histology is the standard for diagnosis and treatment planning and in research settings. Tissue samples provide spatial context of cell types and states into functional tissue units, wherein enriched cell neighborhoods can be indicative of function or dysfunction, especially in a pathological state^1-3^. Understanding these tissue units and neighborhoods bridge the gap between morphological features and the underlying biology. In many cases, histology must be further augmented with specific markers to better understand the underlying pathobiology.

Modern spatial biology studies primarily employ highly multiplexed spatial molecular imaging assays^4-6^. For example, Phenocycler imaging^2^, produces highly multiplexed, single-cell resolution spatial proteomic images for fresh frozen or formalin-fixed paraffin-embedded (FFPE) tissues. While other companion spatial technologies exist^7^, Phenocycler is a broadly used commercial option to achieve the multiplexity required to profile cell types at single-cell or sub-cellular level spatially. A significant challenge shared by many spatial assays is their high barrier to entry for both research and clinical adopters. They require extensive, time-intensive validation, specialized equipment and costly ancillary reagents.

Spatial analysis using digital brightfield histology images using computational methods is an emerging area in biological innovation^8^,^9^. Parallel advancement allows conducting brightfield histology along with spatial molecular mapping in the same tissue sections, thereby creating a one-to-one correlation of molecular information to histopathological information, bridging the molecular and brightfield domains. Capitalizing on this advancement, and to address the limitations for a low resource setup, we implemented a novel convolutional neural network-based semantic segmentation method, DigitAb, for predicting cell types directly using brightfield histology images as input.

Existing segmentation and foundation models involving cell segmentation and classification, such as, CellViT^10^, rely on vision transformer architectures that are limited to small fields of view (typically 224x224 pixels)^11^ affecting accurate segmentation and generalizability. Further, these existing networks are limited to a smaller number of output classes^12^. In contrast, DigitAb incorporates much larger patch sizes for training and prediction (up to at least 769x769 pixels with companion pixel resolution) and is trained on an extensive set of molecular-based cell labels, namely, employing more than 3 million expert annotated kidney cells, using 29 kidney tissue whole slide images (WSIs) spanning multiple subjects and institutions. It also involves domain adaptation end-to-end, and class-balanced training using on-the-fly patch loading and loss calculation for greater generalizability in predicting ten cell types. We achieved a balanced accuracy of 0.78 for kidney histology. As a proof of concept and potential clinical utility, we also show high correlation (*R*^2^ = 0.71) in predicting inflammatory cells for Banff classification of biopsies in kidney transplant rejection, and for association with clinical characteristics in diabetic nephropathy. DigitAb is made available for end-users online, in an easy-to-use system for prediction at scale, and for reproducibility.

## Results

### Overall strategy

To achieve automated segmentation of light microscopy images, our first step was to build a ground truth using network training of paired Phenocycler images **Fig. 1a)**. The DAPI channel from Phenocycler images was used for segmenting nuclei as fiduciaries for cells. This was performed using a combination of deep-learning and classical image analysis methods (**Fig. 1b**). Expression levels were then measured for each protein within each cell (**Fig. 1c**). Segmented nuclei from fluorescence images were also registered to the same-section histological images (**Fig. 1d)**. Mean expression values for each segmented nucleus were used for cell clustering using a Louvain algorithm^13^ (**Fig. 1e**), and cluster identities were mapped to the histological domain using the transformation matrix. Each cluster is assigned a cell label with the assistance of an organ expert. The identity of cell clusters was validated by back-mapping to the fluorescence and histological images (**Fig. 1f**). The annotated clusters were combined into a finalized, histologically linked and molecularly validated cell identity ground truth datasets (**Fig. 1g**). The datasets were then split into training and testing, and the semantic segmentation network was trained using histological patches as inputs, yielding semantic segmentations of cells (**Fig. 1h, i**). The trained model, called DigitAb, was finally used for predicting cell types in external datasets (**Fig. 1j**).

**Figure 1.**
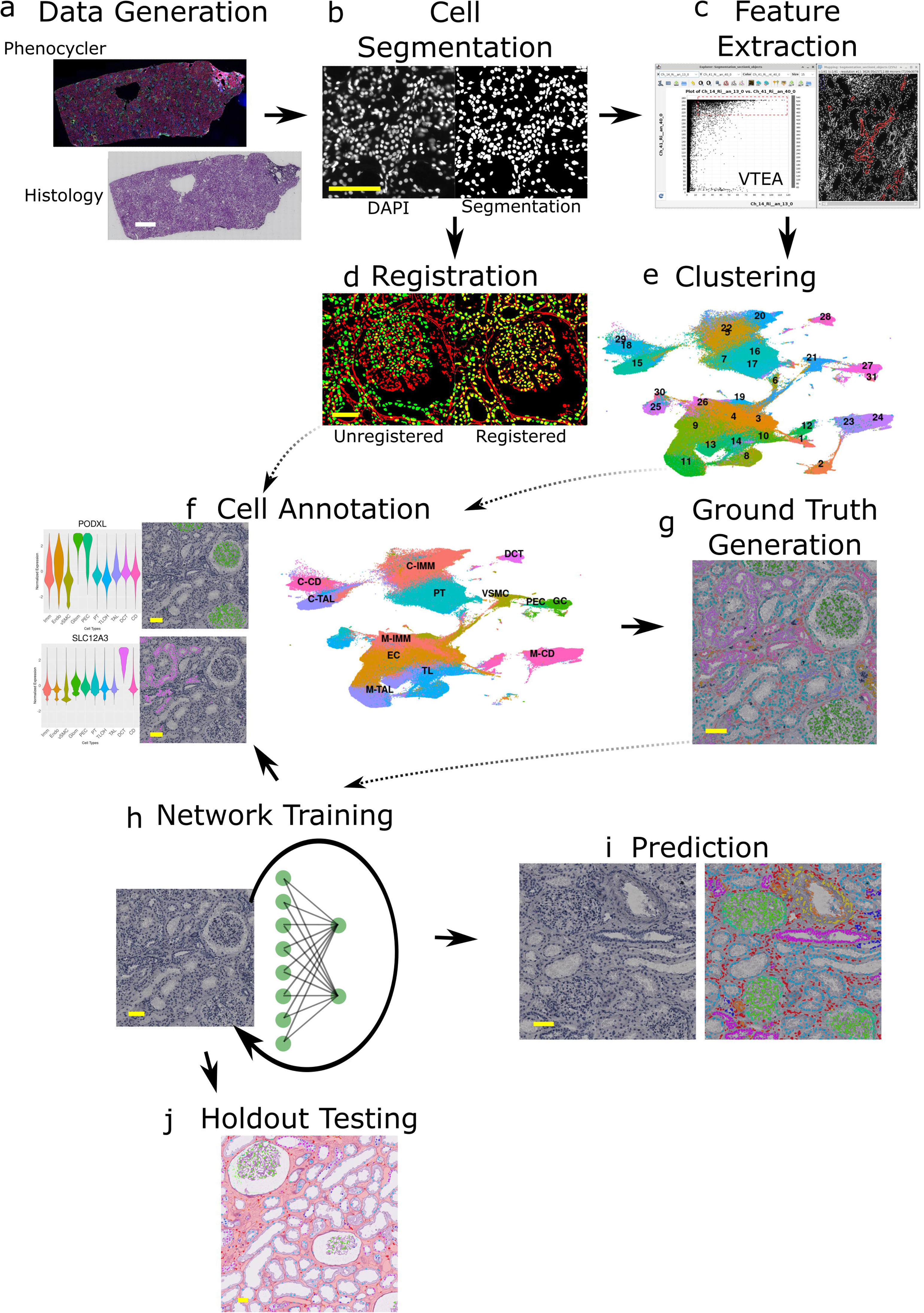
Overview of Cell Labeling and Network Training Pipeline. Given a same-section pair of histology and Phenocycler multiplexed images, segmented cells can be labeled by their proteomic expression and used to train a segmentation network. a) Pairs of histology and Phenocycler images are generated on the same tissue section. b) DAPI images from the Phenocycler image are segmented from thresholding and watershed splitting in the IU-W frozen and IU-K samples, and by DeepCell in the IU-W FFPE samples. Methods were selected by highest mIoU when compared to manual annotations. c) Phenocycler protein expression values are measured from each of the segmented nuclei. d) DAPI segmentations are registered to hematoxylin segmentations using an affine transform. e) Feature matrices are clustered using a graph-based Louvain algorithm. f) Clusters are annotated using canonical cell type markers. g) Clusters are registered to histology to form the semantic ground truth. h) The segmentation network is trained using histology as input, and spatial cell type annotations as ground truth. i) Holdout histology samples are used to test the accuracy of the segmentation network. j) Trained networks are used to predict kidney cell types on external datasets. White scale bars = 2 mm, yellow scale bars = 100 µm.

### Generation of Paired Phenocycler and Histology Datasets across Kidney Cohorts

We first summarize the kidney tissue imaging datasets generated in this study and present representative paired Phenocycler-histology images to illustrate data quality and diversity across cohorts. We assembled multiple kidney tissue datasets spanning multiple preservation and collection methods, each with paired Phenocycler and hematoxylin and eosin (H&E) stained brightfield images (**Fig. 2a-c**). Fresh frozen kidney samples from the Human Biomolecular Atlas Program (HuBMAP) and biopsies from the Kidney Precision Medicine Project (KPMP) were imaged and subsequently post-stained with H&E, yielding paired datasets referred to here as IU-W frozen and IU-K, respectively (**Fig. 2a, c**). The same paired staining and imaging was performed at Indiana University on FFPE kidney sections sourced from Washington University in St. Louis (WUSTL), generating the IU-W FFPE dataset (**Fig. 2b**). Together, these datasets provide representative examples of high-quality, spatially equivalent multiplex immunofluorescence and histology across frozen and FFPE kidney tissues. An overview of sample sources, tissue types, and antibody panel sizes are provided in **Table 1**, while patient demographics and clinical characteristics are listed in **Supplementary Tables 1-3**. A full list of antibody markers is provided in **Supplementary Table 4**.

**Figure 2.**
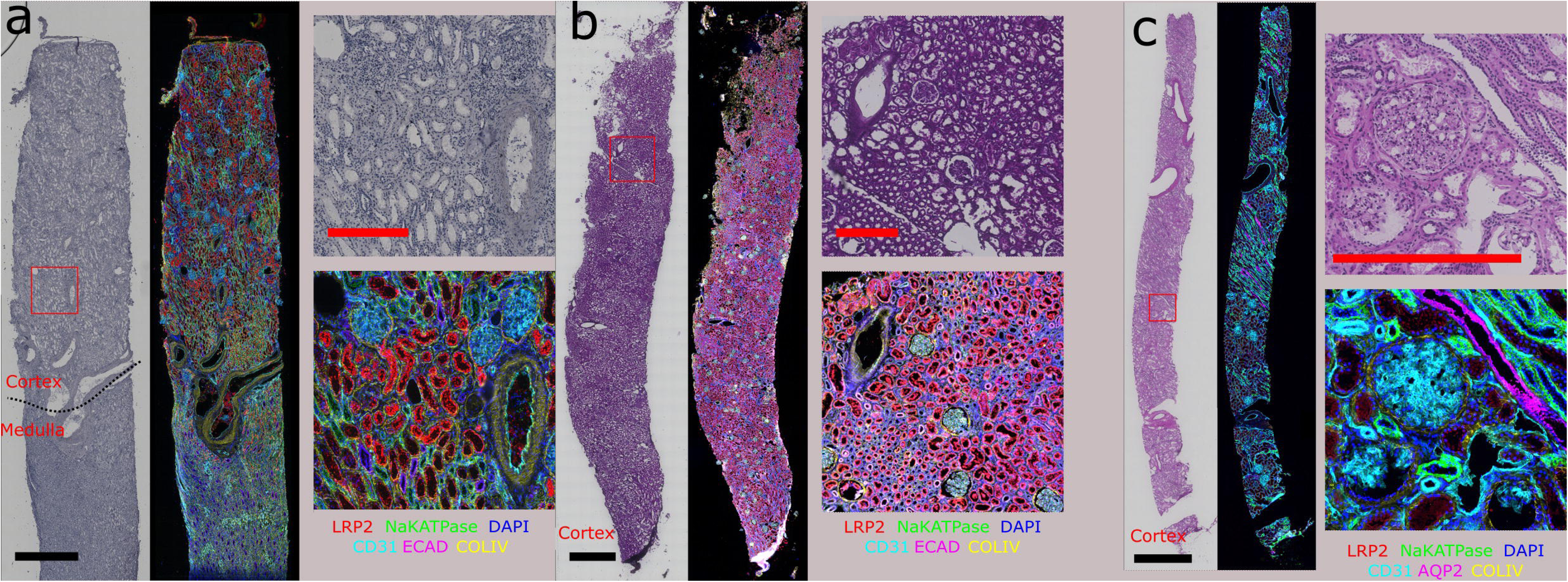
Dataset Generation and Curation from Histology and Omic Pairs. Phenocycler images from Indiana and Washington University frozen and FFPE tissue preparations are post-stained with hematoxylin and eosin. Protein markers shown are labeled beneath Phenocycler crops. a) Fresh frozen nephrectomy samples from sourced from Washington University, imaged at Indiana University. b) Formalin-fixed paraffin-embedded biopsy samples sourced from Washington University, imaged at Indiana University. c) Fresh frozen biopsy samples from the KPMP open-source repository, sourced and imaged at Indiana University. Black scale bars = 2mm, red scale bars = 500µm.

**Table 1.** Sample distribution by dataset. Washington University in St. Louis = WUSTL. Indiana University = IU.

### Ground Truth Generation

To generate ground truth cell maps at a large scale, both molecular and pathomic features of segmented nuclei were used to distinguish cell types in kidney tissue samples.

#### Nuclei Segmentation

Several classical image analysis and deep learning techniques were tested for nuclei segmentation in DAPI, and the top performing methods included local thresholding with watershed splitting in the IU-W frozen and IU-K samples, and a modified DeepCell^14^ segmentation in the IU-W FFPE samples. These two were determined as the best performing methods when quantified against patches of manually annotated nuclei, measured by mean intersection over union (mIoU). Over three million cell nuclei were segmented across all kidney datasets (**Table 2**).

**Table 2.** Number of classified cells by type. Listed for each of the three kidney datasets, cells classified as artifact were discarded.

#### Molecular Feature Extraction

To build a feature set for ground truth cell type classification, summary protein expression statistics were calculated for each segmented nucleus using the open-source volumetric tissue exploration and analysis (VTEA) software^15^. Expression was measured from the segmented nuclear area along with a three-pixel dilated area surrounding the nucleus, to capture cytoplasmic and membrane expression. Markers were classified as nuclear or non-nuclear according to previous studies. See **Supplementary Fig. 1** for Phenocycler expression for various markers overlaid on corresponding histological sections. An additional pair of morphological features were then measured to provide spatial context for each nucleus. These features are i) cortical / medullary classification and ii) distance from the nearest glomerulus. Each feature was measured by comparison with manual annotations of both the cortex, and glomerular boundaries. The extracted features can be visualized in a 2D scatter plot in VTEA, and cells can be gated and highlighted in the corresponding spatial domain. Instructions for running segmentation, feature extraction, cell gating, and other functionalities are outlined in the VTEA documentation (https://vtea.wiki/).

#### Cell Clustering

The final feature set included mean values of protein expression (excluding injury or proliferation markers), and the two compartment-based morphological features. The features were Z-normalized at the patient level, and each dataset was clustered separately to limit batch effect. In the IU-W frozen datasets, 31 clusters were identified using 41 markers for the 2.5M cells, 15 clusters were identified using 46 markers for the 250,000 cells in the IU-K sections, and 19 clusters were identified using 41 markers for the 750,000 cells in the IU-W FFPE samples. A uniform manifold approximation and projection (UMAP) plot for each dataset was constructed to show cluster proximity in low dimensional space (**Fig. 3a**), and violin plots were generated to highlight feature expression across clusters (**Fig. 3b**).

**Figure 3.**
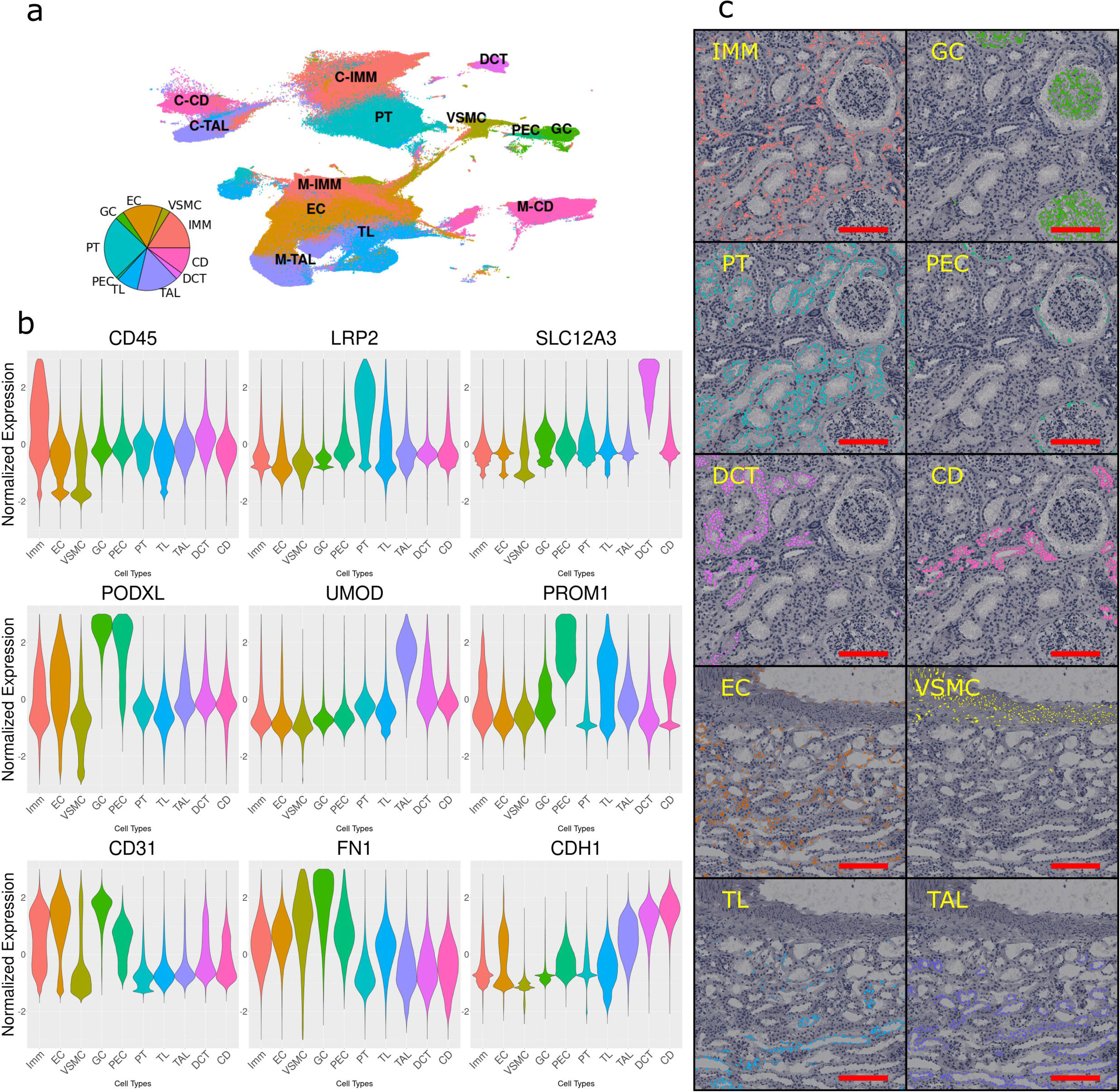
Cell Type Annotation and Ground Truth Generation. Labeling of segmented cell types was completed by Louvain clustering and supervised annotation. a) Uniform manifold approximation and projection (UMAP) plot showing the distribution of cell types in the molecular feature space. b) Shows specific protein expression in several canonical and other discriminative markers. c) Shows the spatial distributions of the clustered and annotated cells. The full list of protein markers used across each panel located in **Supplementary Table 5**. Scale bars = 200µm.

#### Registration of Molecular and Histology Images

Clustered cells were mapped back to segmented cells in the spatial domain (**Fig. 3c**). For registration to histology, nuclei were also segmented from the corresponding H&E sections using color deconvolution of the hematoxylin stain, followed by local thresholding. Registration was performed using an affine transform with the histology segmentation as the reference image, and the DAPI segmentation as the moving image. The transformation matrix was then used to register the cluster maps to their spatial locations. Small adjustments to the hematoxylin segmentation or registration algorithm were made until the structural similarity index was greater than 0.99.

#### Cluster Annotation and Refinement

Classifications of cluster identities were performed in multiple steps with the assistance of domain experts. Starting with a single cluster, the violin plots of protein expression were examined to determine upregulated features, and registered maps were used to determine spatial organization within the full population. Anatomical Structures, Cell Types, and Biomarkers tables from the Human Reference Atlas (HRA)^16,17^ and verification by manual examination of imaging data was done to finalize cell type assignments by the subject matter experts. If a cluster consisted of mixed cell types, it was re-clustered and re-examined with markers, and each sub-cluster was assigned a more granular annotation. For example, *Cluster #*6 in the FFPE dataset harbored a mixture of immune and endothelial cells (**Supplementary Fig. 2**). These cells were then sub-clustered using a reduced feature set of CD31 expression, and a suite of immune markers (e.g. CD45, CD3, CD8). The classification procedure was then repeated for each subclass within the larger cluster population. Another mixed cluster contained endothelial cells and vascular smooth muscle cells within larger blood vessels. These types were separated by thresholding the CD31 expression. Once all clusters and subclusters in a dataset were assigned labels and confirmed with domain experts, clusters with the same label were combined. In some cases, cell distributions and classifications were revised following rounds of network prediction. For example, *Cluster #*8 in the frozen dataset was originally labeled as “medullary pericyte”, but following network predictions, it was found to be morphologically consistent with a thin limb epithelial cell. We built a 10-cell type ground truth, and a finalized class distribution, and the justifications for annotations are listed in **Supplementary Tables 7-9**. The final distribution was the same for all three data sets.

### Network Design

The finalized nuclear maps are used as a ground truth in the training of a semantic segmentation network. A variety of network configurations and pretrained weights were tested first to determine the optimal architecture for training. The most reliable architecture across datasets consisted of a fully convolutional network with a ResNet-50 backbone^18^ and Deeplab V3+ decoder^19^, pretrained with ImageNet^20^ weights (**Fig. 4a**).

**Figure 4.**
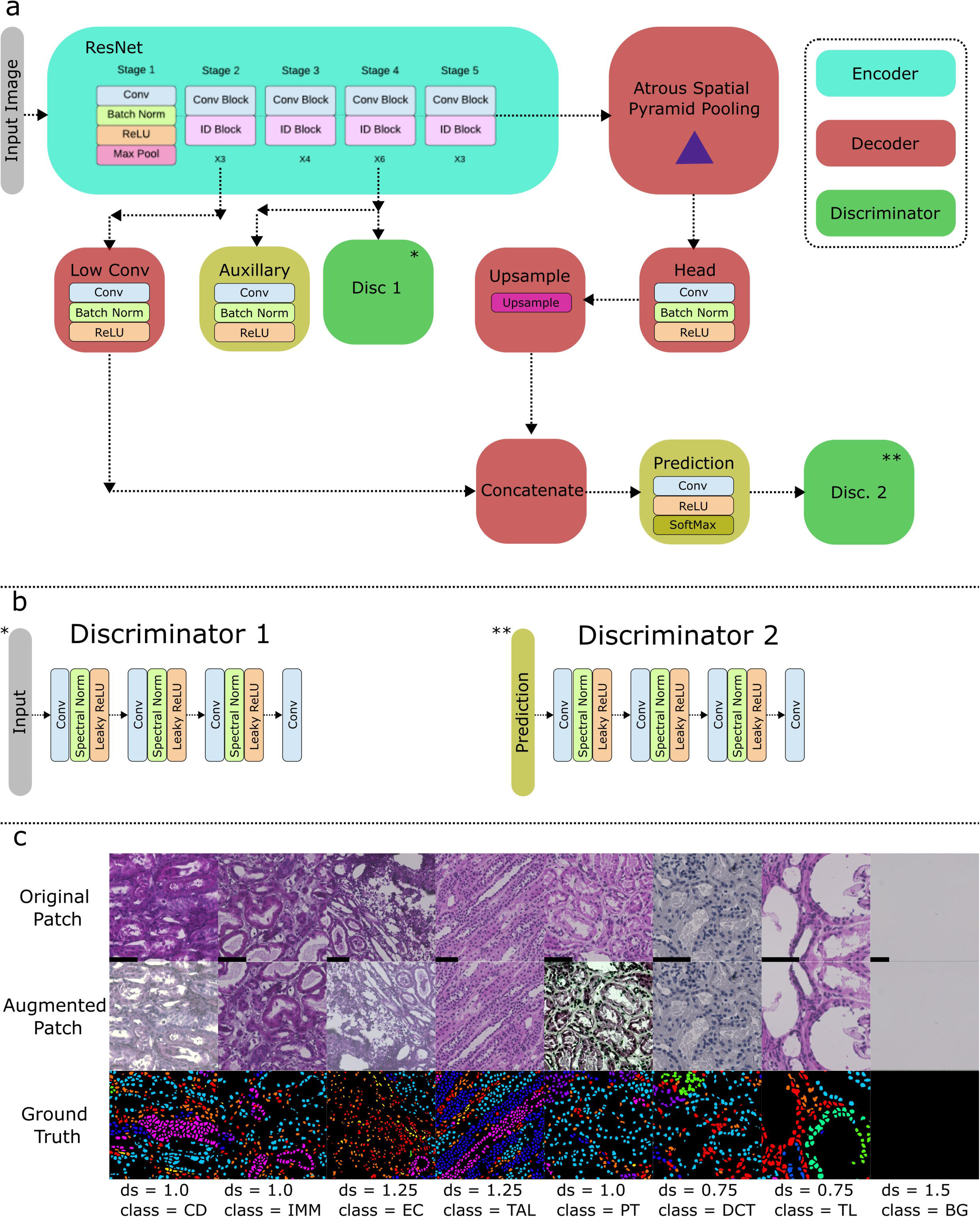
Segmentation Network Architecture. The general encoder-decoder architecture of the segmentation network. a) The image is input to a ResNet-50 encoder. An upsampled atrous spatial pyramid pooling of the highest resolution output is concatenated with a low-resolution output to incorporate information from multiple scales. b) Two discriminators are fit to the encoder and decoder, respectively. Contrastive loss functions assist in producing domain invariant features in the encoder, and realistic segmentation outputs in unlabeled data. c) An example of an eight-image batch for network training. Columns represent a raw image patch, the same patch post augmentation, and a label image. ds = downsample ratio, class = cell type selected by on-the-fly patch loader.

#### Proposed Network & Loss Function

The novel combination of the ResNet encoder and Deeplab V3+ decoder utilizes a lightweight framework with increased depth and combines features at multiple biologically relevant resolutions for both fine and broad representations. An auxiliary supervision branch was added to the latent features via a shallow fully convolutional network with its own cross-entropy loss. This auxiliary loss improves gradient flow, accelerates convergence, and encourages the encoder to learn more discriminative and semantically rich features by providing additional guidance during training^21^. It also acts as a form of regularization, helping to mitigate overfitting and improve generalization. We employed a student-teacher framework where the teacher model is updated as an exponential moving average of the student model’s weights, which smooths out noisy updates and improves generalization and robustness of the learned representations over the course of model training^22^. The network employs a pixel-wise cross-entropy loss function with class weighting.

#### Domain Adaptation

To improve generalization for multi-institutional data, we implemented a novel unsupervised adversarial domain adaptation training strategy, and integrated end-to-end with our proposed semantic set-up (**Fig. 4b**). We here teach the network to extract similar features from similar images in different datasets which are unlabeled and undergo various analytical differences such as thickness, staining, and preparation. Our implementation employs two convolutional discriminator networks attached to the ends of both the encoder and decoder, and a two-step training process is employed. Namely, first the adversarial training combines both supervised cross-entropy loss with adversarial loss, providing the discriminators with incorrect labels for the target images (unlabeled, external). This step measures how well the encoder and decoder can fool the discriminators by extracting domain invariant features, and realistic segmentation masks, respectively. Next, the discriminators are trained to recognize the domain each image is coming from by providing them with the correct labels for each image. This process is designed to slowly improve the feature extraction and decoding by using the unlabeled images during training, by updating the weights of the discriminators at the same time. Adversarial loss weighting is tuned with a hyperparameter, λ, to balance adversarial training with supervised segmentation training.

#### Network Hyperparameters & On-the-Fly Patch Loading

The network was trained with a batch size of 8 and a patch width of 769 pixels. To save disk space and sample the training space more efficiently, patches were loaded *on-the-fly* from our previously published custom patch loader^23^. To load a patch, a random class is selected from the ground truth distribution, and a random image patch from the WSI containing that class is loaded (without pre-patching the WSI). This novel training strategy oversamples minority classes to balance the training. The loader also chooses a random scale factor from a user-defined list, to upsample or downsample patches for multi-resolution training. This training strategy is beneficial for training on WSIs scanned at different magnifications and learning spatial relationships at various microscopic scales. During training the patches are also run through a series of random augmentations. An example training batch is shown in **Fig. 4c**.

### Segmentation & Classification of Kidney Cell Types

#### Network Evaluation

Cross-validation was implemented to test DigitAb performance for individual institutes. Samples from a single patient were held out for a single fold and used for model testing and validation. WSIs were predicted using an overlapping moving window approach, and then stitching predicted patches together by removing an outer ring of pixels of user-defined size. This stitching is designed to mitigate edge artifacts. Due to the vast prevalence of background pixels, thus inflating cell-type specificity, only pixel-wise sensitivities for each cell type were quantitated. Sensitivity was then averaged over each cell type to treat each class equally, regardless of prevalence.

#### Frozen Samples

In this experiment, only samples from the HuBMAP IU-W frozen dataset were used for training and testing. As there were five patients in this dataset, the test results were averaged over each fold. The network receives a histology image patch (**Fig. 5a**) and outputs a class-wise map, which is then overlaid with the original histological image (**Fig. 5b**), and compared to the ground truth (**Fig. 5c**). Our collection of trained models achieved a balanced accuracy (mean sensitivity) of 0.78 in ten cell types (**Fig. 5d**), a 0.68 improvement over a 0.10 random baseline. SoftMax outputs of the segmentation network were also output to measure the confidence of the network in each cell classification. All cell classes achieved an area under the curve (AUC) greater than 0.9 (**Fig. 5e**). Herein no domain adaptation was implemented.

**Figure 5.**
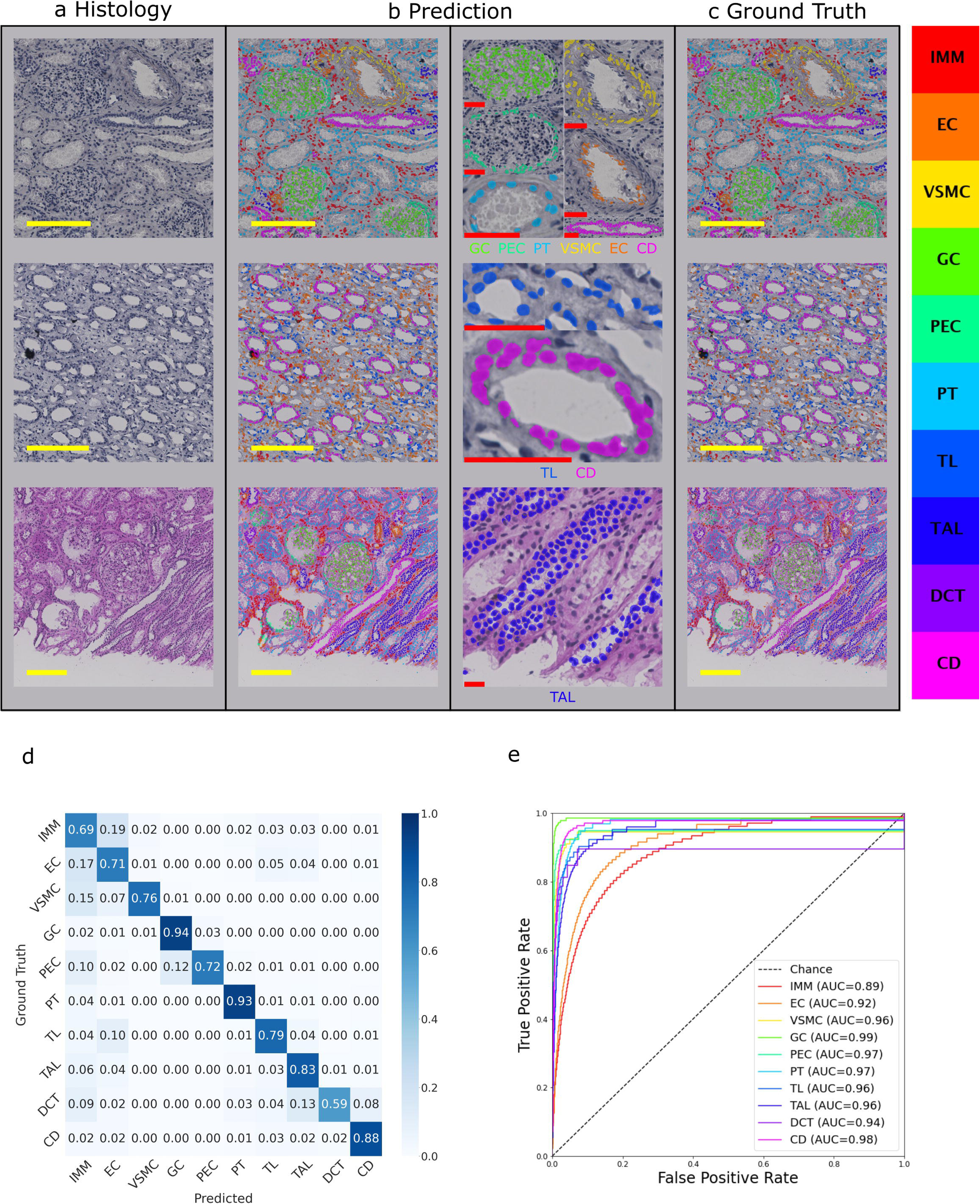
Segmentation Network Performance in IU-W Frozen Dataset. Network predictions are shown in holdout samples during cross validation. a) shows native histology images in the fresh frozen cortex and medulla (rows 1-2) from nephrectomy tissue and from needle core biopsy (third row) cohorts. b) colored overlays for cell types predicted by models during cross validation. c) Colored overlays for ground truth cell types. d) Confusion matrix for full cross-validation results in the fresh frozen cohort, showing pixel-wise classifications by the network vs our ground truth. Box colors are normalized by row sum. e) Receiver-operator characteristic (ROC) curves for each cell type. Cell type key is shown in colorbar. (IMM = immune cell, EC = endothelial cell, VSMC = smooth muscle cell, GC = glomerular cell, PEC = parietal epithelial cell, PT = proximal tubule epithelial cell, TL = thin limb epithelial cell, TAL = thick ascending limb epithelial cell, DCT = distal convoluted tubule epithelial cell, CD = collecting duct epithelial cell). Yellow scale bars = 200µm, red scale bars = 50µm.

We observed classification confusions between the thick ascending limb (TAL), distal convoluted tubule (DCT), and collecting duct (CD) epithelial cells of the distal nephron. TAL and DCT cell types have specific markers for classification, are in proximity along the tubular network, and present similar morphologies in the brightfield images, while markers used for CD to generate ground-truth were general epithelial markers (CHD1) and present more morphologically distinct than the former. Another network confusion was observed between immune and endothelial cells. We found endothelial cells in smaller vessels like peritubular capillaries were often confused as interstitial immune cells. This is likely due to the similar spatial niche, and the thickness of the sections which may obscure the peritubular capillary lumen that would otherwise be visible in a thinner section. Conversely, the performance of endothelial cell prediction was higher in the medulla as opposed to the cortex. This might suggest that the cells of the vasa recta and other medullary vessels may be more easily identifiable in thicker sections. This may also be explained by the larger distance between tissue structures in the medulla.

#### FFPE Samples

Samples from the HuBMAP IU-W FFPE dataset were used for training and testing. Each WSI came from a unique patient, so each WSI was held out for a single fold and used for testing. Our collection of models achieved a balanced accuracy of 0.69 across 9 cell types (**Supplementary Fig. 3a**), a 0.58 improvement over baseline. No domain adaptation was implemented herein.

Since these sections did not contain any medulla, there were no thin limb epithelial cells annotated in the ground truth. This may explain the drop in performance in endothelial cells, since as previously mentioned, endothelial cell performance was higher in medullary regions. We observe a reduced segmentation performance in the IU-W FFPE samples, despite the same preprocessing and training procedures. Notably, the IU-W FFPE dataset was smaller in sample size and tissue per image and exhibited great heterogeneity across samples. We also observed network confusions in the distal nephron, similarly to the frozen dataset results. However, predictions in both proximal tubule and glomerular cells measured 0.94 sensitivity.

### External Testing and Visualization

#### Reference Kidney from KPMP

We tested generalizability of DigitAb using datasets external to the IU HuBMAP (KPMP) and combining the previously discussed IU-W frozen and FFPE labeled samples for training. The studies herein were conducted with and without domain adaptation.

The IU-K healthy reference tissue living donor biopsy data was used. The samples contain both Phenocycler and histology images, and were interrogated with a nearly identical marker set as in the training, yielding same ground truth labels. A single DigitAb model was trained using all samples from both the IU-W frozen and FFPE datasets. The network achieved a balanced accuracy of 0.54 in 10 cell types (**Supplementary Fig. 3b**). The distal nephron epithelial cell confusion is found to be even more apparent in this dataset, which may again be attributed to minimal medullary tissue.

To assess whether differences in model performance across datasets could be attributed to histological batch effects, we performed HistoQC^24^ analysis on the three datasets. Feature-based clustering revealed clear separation between datasets, indicating the presence of a batch effect arising from differences in tissue sourcing, preservation, and staining (**Supplementary Fig. 4**). While the IU-K tissue is fresh frozen, these samples are patient biopsies limited to the cortex, whose stain distribution is visually distinct from both the IU-W frozen and FFPE samples.

### Application to Unseen and Untested Clinical Biopsies

To demonstrate DigitAb’s clinical utility and broad usability, we used biopsies from kidney transplant patients to assess inflammation and diabetic nephropathy kidney biopsies to examine impact of inflammation on kidney function decline. We trained a model using IU-W frozen and FFPE and IU-K samples as labeled training, with the diabetic samples as unlabeled training samples. Using the resulting optimized domain-adapted model, these samples were then predicted for cell type annotation. For transplant biopsies, we examined time-zero (t0), baseline, and for cause periodic acid-schiff (PAS) stained kidney tissue biopsy WSIs^25^. The WSIs were segmented for functional tissue units (FTUs) by our earlier published multicompartment segmentation method^26^, and annotated for cell types using domain-adaptive DigitAb. The multicompartment segmentation algorithm segments the cortical and medullary interstitium, glomeruli, tubules, and arteries/arterioles. We focused on DigitAb’s immune cell prediction for inflammation quantitation (**Fig. 6a-e**). For clinical significance, across 19 WSIs (one per patient), a total of 1.8M nuclei from 313 mm^2^ of tissue were annotated, with a total computation time of 44 minutes and 23 seconds.

**Figure 6.**
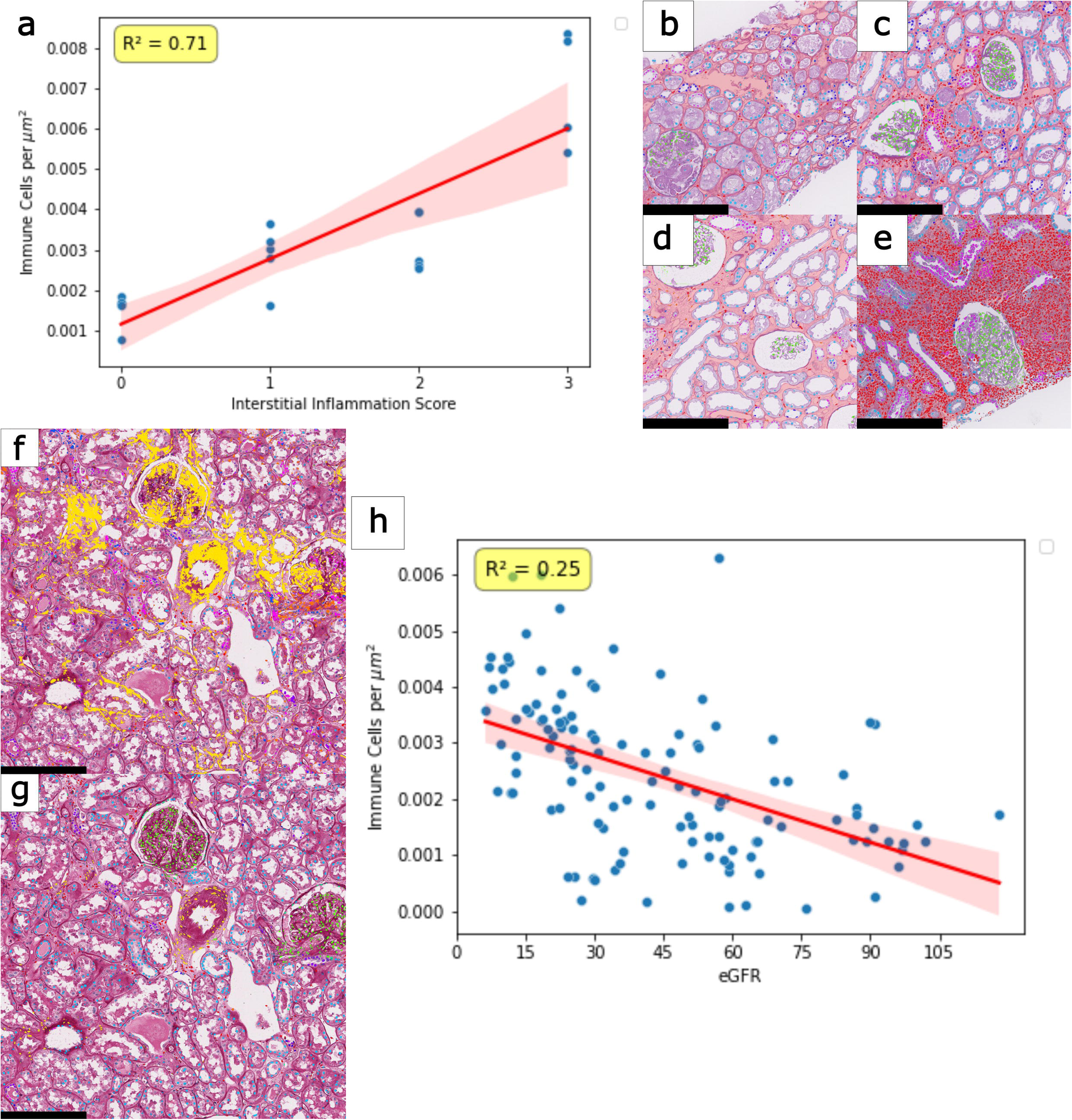
Segmentation Applications in External Datasets. a) Correlation of immune cell count density in the cortical interstitium, with the pathologist-adjudicated Banff inflammation score. Examples shown in inflammation cases with scores 0-3, represented by b)-e), respectively. Histology section from DN cohort, with overlaid predictions for f) non-domain-adapted DigitAb model, and g) domain-adapted DigitAb model. h) Correlation of immune cell count density in the cortical interstitium, with baseline eGFR. Black scale bars = 250µm.

Full tissue area, cortical interstitial area, immune cell area, and immune cell count were measured from the resulting FTU segmentations and cell type annotations. From the segmented areas, we calculated the immune cell count densities in the cortical interstitium and compared these values against the gold-standard – pathologists examined and quantitated Banff inflammation scores^27^. Using linear regression, we found a coefficient of determination of 0.71 (**Fig. 6a**) between computationally derived vs Banff inflammation scores, suggesting fair concordance with pathologists. Hence, in completely unseen clinical samples, DigitAb can assess inflammation and resolve according to clinical criteria and inform on quantitative metrics that would be otherwise laborious. DigitAb is also able to evaluate the proximity of immune cells to different FTUs, as FTU-immune neighborhood can inform on the underlying nephron segments that are injured and signaling mechanisms^28,29^. For example, DigitAb analysis shows greater percentage of immune cells aggregate near distal nephron tubular segments in the cortex as inflammation severity increases. By binning counts of immune cells into their nearest FTU, we compared patients across inflammation scores using Kendall’s rank correlation coefficient (τ)^30^ (**Table 3**.). Two FTU bins showed statistically significant associations with increasing inflammation severity. The collecting duct bin showed a strong positive association with inflammation severity (τ = 0.60, *p* < 0.001), while the proximal tubule bin showed a strong negative association with inflammation severity (τ = -0.63, *p* < 0.001). Associations among the other FTUs were not statistically significant. This indicates that increases in inflammation severity are associated with selective redistribution or recruitment of immune cells towards the distal end of the nephron. This observation would need further testing and confirmation in their impact on renal injury and recovery.

**Table 3.** Kendall Tau coefficients by Functional Tissue Unit. Coefficient explains magnitude and direction of association with increasing inflammation severity in transplant biopsies.

#### Clinical Association in Diabetic Nephropathy

We next examined the performance of DigitAb on PAS-stained kidney FFPE biopsies from patients with diabetic nephropathy. We annotated 13.5M nuclei from 135 WSIs with a total tissue area of ∼4,014mm^2^, requiring 9h 45m of computation. Segmentation was repeated for the IU-W frozen sample trained model, to qualitatively compare how domain adaptation improves model performance. Before domain adaptation, cell type predictions heavily favor smooth muscle cell predictions (**Fig. 6f**), and do not follow the contours of the nuclei. While post domain adaptation, cell type predictions are more balanced and are more representative of the actual cell contours (**Fig. 6g**). Additionally, the WSIs were segmented for FTUs by our earlier published multicompartment segmentation method^26^, and the density of immune cells was calculated for each sample. These metrics were then associated with estimated glomerular filtration rate (eGFR) (**Fig. 6h**). We measured *R*^2^ = 0.25 between immune cell density and eGFR, where a higher density was associated with a lower eGFR. Using follow-up measurements of SCr, we calculated a 3-year composite outcome for each patient, where a progressor was defined as an individual whose SCr doubled, eGFR declined by 50%, or who reached end stage kidney disease within that 3-year period. We repeated the previously discussed proximity study with this dataset and performed Mann-Whitney U tests for each FTU bin. However, no single bin showed a statistically significant difference in patient-level proportions (all *p* > 0.20). The collecting duct did show the highest magnitude in median difference between outcomes, but this was not found to be statistically significant. Despite the lack of FTU-specific differences, a permutation-based global composition test indicated a statistically significant difference in the overall distribution of immune cells in the samples (*p* < 0.05). This does suggest a difference in immune cell proximity when considered jointly. A simulation-based power analysis demonstrated that the observed global shift in bin-level composition is underpowered at our sample sizes. Conventional power (> 0.8) was achieved when simulated sample sizes reached 150 patients per outcome group. This indicates that the observed effect is too diffuse to be reliably detected in our cohort.

### Accessibility and Reproducibility

Models and select data for testing with DigitAb generated outputs are available for end-users via cloud using our CompRePS platform^31^ at *athena.rc.ufl.edu*. Cell segmentation and classification, especially, for kidney histology images can be conducted by end-users using our cloud system, with a single click, and in an easy-to-use fashion. Information and figures for navigating and running the networks are outlined in **Supplementary Information**. Network predictions can be output as tiff images, xml, json, and geojson formats. This availability of data and model establishes reproducibility of our results and allows researchers from any setup to employ DigitAb to generate high throughput cell annotation data for kidney histology.

## Discussion

Typical renal biopsies contain tens of thousands of cells spanning multiple distinct cell types, making detailed manual labeling prohibitively time-consuming and susceptible to inter-observer variability. We developed DigitAb for automated granular annotation of digital histopathology WSIs, enabling more precise spatial and cellular analyses, ultimately improving the interpretability of tissue architecture, inflammation, disease processes.

DigitAb leverages a rich but limited molecular datasets of Phenocycler and same section histology to teach an AI network to infer specific cell types directly from conventional histology. By transferring molecular information to H&E images, the model enables similar levels of cellular representation without requiring specialized molecular imaging. Importantly, the tool demonstrates concordance with pathologist estimations of inflammation in renal transplant biopsies, association with clinical characteristics, and allows for quantification in the distribution of immune cells across various patient populations, underscoring its potential clinical utility. Online availability removes barriers related to local infrastructure, facilitating broader adoption and reproducibility of our results. DigitAb could also be used for augmenting pathologists’ efforts.

DigitAb is implemented using various computational novelties, including its modular network architecture, supporting customization for diverse tissue types and segmentation tasks, on-the-fly patch loading and custom loss function design for efficient computation as well as increasing the effective sample size through continuous whole-slide sampling, and balancing rare cell types, and patch-level resizing and augmentation, allowing the model to operate across multiple resolutions. Domain adaptation strategies mitigate the limited training data inherent to expensive molecular modalities.

Despite these strengths, DigitAb also has several limitations in its current form. The comparatively lower performance observed in IU-W FFPE and IU-K is likely driven by a combination of sample size and domain-specific image characteristics. The IU-W frozen samples exhibit a wider range of tissue structures spanning from the renal capsule to the inner medulla, while the former datasets are limited to smaller areas of just the cortex. This limited the model’s exposure to a broader distribution of learnable tissue structures, where a larger number of training samples and increased model capacity would be needed to improve generalization. This likely reflects a data-specific limitation rather than a failure of the modeling approach. HistoQC analysis demonstrated the distinct phenotypes of each labeled dataset, and are expected, given the differences in tissue preservation and processing.

As a more general limitation, Phenocycler imaging requires histologic sections thicker than those routinely used in diagnostic practice, reducing the visual detail of the images and limiting the information learned by the model. The cyclic imaging process itself may also contribute to tissue degradation. Tissue sourced for this project were also classified as healthy reference, limiting the exposure of the model to pathological lesions. Extending the network to diseased samples is difficult in severely affected tissue, since canonical markers of various cell types may diminish in injured states, thus making it difficult to classify accurately the ground truth. Molecular markers used for cell type identification are also not strictly a binary problem and often exist along a continuous gradient, introduce ambiguity often, leading misclassifications in cell annotations. These biological gradients may impose an upper bound on the achievable accuracy of performance assessments. Furthermore, training data are derived primarily from H&E-stained sections, whereas renal pathologists additionally use PAS and other special stains. Training on additional stains and integrating these into the model will improve utility. Finally, evaluating generalizability remains challenging because obtaining accurate labels for standard whole-slide images is difficult, limiting large-scale external validation.

## Methods

### Institutional Review Board Approval

This study was approved by the institutional review boards at the University of Florida (IRB202201030). All human tissue samples analyzed were obtained under IRB-approved protocols from contributing institutions. Samples were fully de-identified prior to access and analysis. Informed consent was obtained by the contributing institutions, and no compensation was provided to participants.

### Data

#### The HuBMAP Consortium^32^

Phenocycler and associated histology imaging data were sourced from patients and generated at the IU School of Medicine. Fourteen Phenocycler and hematoxylin and eosin (H&E) frozen sample image pairs from five subjects were generated according to published protocols^33,34^. First, the tissue is cyclically tagged and imaged with a total of 40 protein markers (not including DAPI). These markers were specifically chosen to highlight and discern the various major cell types in the human kidney, following the guidelines of the human reference atlas (HRA)^16,17^. These marker names and their targets are listed in **Supplementary Table 4**. Following the Phenocycler workflow, the same tissue sections are stained with H&E and scanned as digital images at 20X magnification.

An additional five Phenocycler and H&E stained FFPE image pairs from five subjects were generated according to established protocols^33,34^. These samples are tagged and imaged with a slightly different but largely overlapping collection of 40 protein markers. These samples are also stained with H&E and scanned as digital images at 20X magnification, at 0.5 μm/pixel resolution.

#### The KPMP Consortium

External data was sourced from the KPMP healthy reference tissue (HRT) repository. The KPMP data consists of eight frozen tissue samples imaged with 45 molecular markers (not including DAPI), and corresponding same-section H&E-stained histology imaged at 20X magnification, at 0.5μm/pixel resolution.

#### The University of California at Davis (UCD) Transplant Rejection Cohort

Samples were also sourced from the pathology archives of UCD. Nineteen baseline, time-zero (t0), and for cause transplant biopsies were used for this study. Tissue sections were prepared at 2-3 µm thickness and stained with Periodic acid-Schiff (PAS). Slides were scanned using a brightfield microscopy whole-slide scanner and at 40X magnification, at a 0.25 µm/pixel resolution. WSIs were graded according to Banff criteria for rejection (**Supplementary Table 5**).

#### The Washington University in St. Louis (WUSTL) Diabetic Nephropathy Cohort

Kidney biopsies from healthy control patients, and patients with diabetic nephropathy were obtained from the pathology archives of Washington University Kidney Translational Research Center. Slides were scanned using a brightfield microscopy whole-slide scanner (Aperio ScanScope, Leica Biosystems, Vista, CA) at 40X magnification and 0.25 µm/pixel resolution. Patient metadata included age, sex, race, estimated glomerular filtration rate (eGFR), and serum creatinine (SCr). Also included are follow-ups for end stage kidney disease, doubling of serum creatinine, and 50% decline in eGFR (**Supplementary Table 6**).

### Quality Control and Batch Identification with HistoQC

The open-source software HistoQC^24^ was used to analyze the low-level image features in the histology samples from each of the IU-W frozen, FFPE, and IU-K datasets. The existing “first” configuration was used to run the initial analysis. The open-source CohortFinder^35^ software was then run on the same collection, using the number of clusters set to -1, to let the built in algorithm fit the data to an optimal number of clusters.

### Nuclei Segmentation

Segmentation of the DAPI in Phenocycler images was completed empirically by two methods. Namely, several 512x512 pixel crops of the DAPI channel from a few sections were manually annotated as ground truth, of which segmentation performance was quantified and compared against, and the best method was selected for each preparation (frozen vs FFPE). For each image, histogram equalization is used to enhance the contrast. The first segmentation method uses the *Connect 2D/3D with kDTree* method in VTEA^15^, where Otsu’s thresholding is used to separate foreground from background, and watershed post-processing is used to separate overlapping nuclei. The second segmentation method is a combined deep learning and classical image analysis approach. Namely, DeepCell API^14^ is used to segment the contrast enhanced nuclei, which automatically performs watershed post-processing to split overlapping nuclei. Since the WSIs are too large to segment at once, patches of 1024x1024 pixels are passed to the API, and segmented images are eroded with one pixels to prevent neighboring objects from merging in post-processing. Border crops of 128 pixels were removed to prevent edge artifacts.

### Molecular Feature Extraction using Volumetric Tissue Exploration and Analysis (VTEA) Software

Spatial analysis and feature extraction is performed in the open-source VTEA software^15^. Using the segmentations, molecular features are extracted from each cell. Since some of the molecular targets are in the cell cytoplasm or membrane, a ring dilation distance of three pixels around the nuclear segmentations is used to capture additional cytoplasmic/membrane features. From the corresponding spatial locations where cells were segmented in the DAPI channel, mean values, minimum/maximum values, and standard deviations are measured from each of the other Phenocycler channels. These values are measured for both the nuclear and cytoplasmic representations.

To provide spatial context for cell localization within the kidney, two additional morphological features are extracted from each nucleus. First, the cortex is manually annotated, and cells are assigned as cortical or medullary to better separate cells specific to each section of the kidney (i.e., thin limb/loop of Henle in medulla, or distal convoluted tubule in cortex). Second, each of the glomeruli in the samples are manually annotated, and a map of distances to the nearest glomeruli are measured for each nucleus. This distance feature is used to assist in the classification of glomerular cells and as an additional metric for cortical/medullary orientation.

#### Clustering

Cell by feature matrices for each of the images are then exported from the VTEA software, the morphological features are concatenated, and the full matrix is imported into R. Features are first log-normalized and z-normalized to limit batch effect across samples within a dataset and then concatenated into a single large matrix. These feature matrices are analyzed separately for different tissue preparations, due to the differences in Phenocycler marker panels and respective signals. For clustering, injury and proliferation (cell state) markers are omitted. The FastPG package^36^ employs a computationally and time efficient Louvain-based clustering method, and is used to partition the cells into cluster with similar feature values.

Violin plots are generated to visualize the expression of each Phenocycler marker and morphological feature value in each of the clusters, and to summarize all the feature values across a single cluster. The cluster values are then mapped back to the corresponding nuclear segmentations, for spatial visualization.

The nuclei in the corresponding H&E sections are then segmented by deconvolving the H&E image into a single channel representing hematoxylin expression, and binarizing with an Otsu’s threshold^37^. This is done to provide a reference image for registration of the cluster map, which is done using an affine transformation.

#### Cell Identity Mapping

The measured feature values, and spatial information in both the Phenocycler and histological domains assist in assigning cell-type labels to each discovered cluster. Assigning ontological labels is a multi-step process involving several domain experts. First, a Uniform Manifold Approximation and Projection (UMAP)^38^ plot is generated to project the feature values for each cell into a 2D plot, with cluster values used to color the points. Then, each cluster is interrogated from one ordinal direction to the other, considering the proximity of other clusters within the plot (e.g. epithelial subtypes will likely cluster near each other). The profile of the selected cluster is examined, determining which Phenocycler markers or features may be positively expressed or absent. Based on the combination of upregulated/down-regulated features and domain knowledge in spatial biology, an assignment is suggested for the cluster. Then, the histological morphology and co-localization of the cluster is examined to confirm or alter the label. If a cluster is deemed to contain more than one cell type label (e.g. endothelial and smooth muscle cells in small vessels), the clustering process is repeated just within the current cluster, and labels are then assigned for each subcluster. Features are selected based on the cell types included in the larger cluster. For example, for a cluster containing multiple types of epithelial cells, only epithelial markers and morphological features will be used for subclustering. Following this process, multiple clusters with the same labels are combined, and a cell class distribution is finalized. The same transformation matrix used for registration is then used to register the cell map to the histological domain. Justifications for cluster annotation are outlined in **Supplementary Tables 7-9.**

### Neural Network Construction & Evaluation

#### Hardware Specifications

Computational experiments were conducted on the HiPerGator-AI supercomputer at the University of Florida. Each network was trained on 1-Nvidia DGX B200 GPU with 48GB RAM, and 8-CPU cores with 8GB RAM each.

#### Architecture

The architecture selected for this work is a standard fully convolutional network^39^, consisting of a ResNet^18^ backbone with a Deeplab V3+ decoder^19^. The ResNet architecture is chosen to keep the encoder lightweight, while still extracting deep features and without losing performance^40-43^. The Deeplab V3+ decoder is chosen because it can incorporate features at multiple scales, which is useful in a histological context, since both fine details and larger-scale neighborhoods are important for determining cell types.

#### On-the-Fly Patch Loading

During training, a novel on-the-fly patch processing is used to expedite the training, and to oversample minority classes^23^. The patch loader builds training batches by directly sampling from the WSIs. This is done by providing a text file list of WSIs to be used during training, and a random slide is selected from this list. The loader then selects a random class from the provided ground truth, and then finds a random place in the WSI where the selected class is present and centers a patch around the object. The patch is then randomly shifted by less than or equal to half the patch size in the horizontal and vertical directions. Lastly, the patches undergo a series of randomized augmentations according to user defined probabilities, including random flipping, resizing, color mapping within a range of LAB color space^44^ parameters, and gaussian blurring. Each patch also undergoes “hard” augmentation with a low probability, which includes a random selection of contrast augmentation, histogram equalization, inversion, blurring, brightness augmentation, hue shift, sharpness augmentation, posterization, and solarization. This process repeats in parallel for all the patches in a full batch.

#### Hyperparameters and Supervised Loss

Network hyperparameters were optimized using RayTune^45^, which utilizes Bayesian optimization for parameter search and selection. RayTune also uses early stopping to speed up the process in cases of suboptimal performance with a particular combination of parameters.

For the ResNet-based model, an online hard-example mining (OHEM) cross entropy loss function is employed to focus training more on hard to classify samples^46^. Stochastic gradient descent is used for optimization, with momentum = 0.9, and an initial learning rate of 0.002, with a polynomial decay of power = 0.9. A batch size of eight, 769x769 pixel patches is used, and validation loss and performance metrics were measured every 100 training steps. A fully convolutional auxiliary module is also attached to the end of the encoder to improve gradient flow and encourage discriminative feature extraction.

#### Class Structure Optimization by Backward Elimination

During preliminary experiments, a highly granular and extensive cell class distribution was used to test segmentation. However, many of the highly granular and rare cell types were being predicted extremely poorly in holdout sections. Therefore, over several experimental iterations, highly specific cell classes were combined into broader categories to improve segmentation without compromising cell class identities. For example, in the initial training, we had cell type labels for both “descending thin limb epithelial cells” and “ascending thin limb epithelial cells”. After multiple rounds of training, we found that nearly all the false positives for one thin limb class would fall within the other thin limb class. These two segments are nearly identical both molecularly and morphologically, and therefore these classes were combined in the next training iteration, since the network could not visually distinguish these two subtypes from only the histological image. The final class distribution consists of 10 cell types, outlined in **Supplementary Table 10.**

#### Network Evaluation

Slides are predicted by first obtaining a mask of the tissue, to eliminate erroneous prediction on patches only containing background. This is done by converting to the hue, saturation, and value (HSV) color space, and thresholding the saturation channel. Based on user-defined size and step values, the slide is partitioned, and the coordinate values of patches are saved in memory. If a patch contains >99% background, the coordinates are deleted from prediction. Slide patches with coordinates corresponding to tissue section are then loaded in parallel, run through the prediction network, and saved in a larger grid. Prediction images are then saved as tiff file with the same x-y dimensions as the histology image. Since the prediction images are the same size as the ground truth, pixel-wise comparisons can be performed using binary logic.

### Prediction Model Optimization for IU Data

In the first experiment, only the frozen sections are used for training, validation and testing. Due to the relatively small dataset size, 5-fold cross validation is used, where all the slides from each of the 5 subjects are held out once in each fold. One slide from each of the training subjects is used as validation, and the network weights are saved each time the validation accuracy reaches a new best. The network is trained for 200 epochs, at 100 steps per epoch. The best model, measured by highest validation accuracy, is used for holdout testing.

WSIs were annotated for any existing artifacts that may influence performance metrics. These included tissue tears, folds, blurry areas, air bubbles, debris blocking the field of view, or staining inconsistencies. The artifact annotations are made available for viewing in CompRePS^31^.

Each of the five models were only evaluated on the slides on which they were not trained. Following predictions, areas of annotated artifact were excluded-Pixel-wise sensitivities were measured for each cell type, and the full results presented are cumulative over each fold.

In the second experiment, only the FFPE sections are used for training, validation, and testing. Five-fold cross-validation is used once again, where each slide is held out once in each fold. The full results presented are cumulative over each fold.

### Model Generalization

#### Domain Adaptation

An unsupervised adversarial domain adaptation strategy is implemented to improve generalization of the network by utilizing unlabeled data during training^47,48^. The general framework has been adapted from a previous publication^39^. Two fully convolutional discriminators are attached to the ends of both the encoder and decoder networks, where the role of the discriminator is to classify whether an image patch belongs to the labeled source domain, or the unlabeled target domain. Both discriminators utilize a binary cross-entropy loss function. A single training step now consists of two phases: adversarial and discriminator training. In the adversarial training phase, the unlabeled target images are fed into the segmentation network. The extracted features are used as the input for the first discriminator, and the SoftMax output of the decoder is used as the input for the second discriminator. The discriminator loss functions are given the incorrect label (the label for source images) to test how well the network can fool the discriminators. The gradients flow through a gradient reversal layer with an adjustable weight parameter^49^. To fool the discriminators, the encoder must not only extract useful features for segmentation, but also features that are domain invariant, while the decoder must produce segmentations that are realistic. The same supervised segmentation loss function is also added in this step. In the discriminator training phase, both labeled source images and unlabeled target images are fed into the segmentation network. This time, the discriminator loss functions are given the correct labels for each image, to teach the discriminators what each domain looks like. The two-stage training is designed to improve the encoder’s ability to extract domain invariant features, and the decoder’s ability to produce realistic segmentations, while also improving the discriminators’ abilities to distinguish the two domains.

#### External Testing

For testing the IU-K dataset, the entirety of the frozen and FFPE datasets were used as training slides. Network training was completed for 200 epochs at 100 steps per epoch, with eight images in each training batch. Network performance was measured and averaged across each of the slides. Performance metrics were kept the same as the first segmentation experiments.

### Use-Case 1: Banff Inflammation Estimation in Transplant Rejection

Baseline, time zero, and for-cause biopsies from transplanted kidneys were obtained from the pathology archives of UCD. These slides were graded on Banff^27^ criteria by a pathologist as part of standard clinical workflow. Nineteen PAS-stained slides were selected with nearly even distribution of inflammation Banff scores. Five slides from Grade 0, five slides from Grade 1, five slides from Grade 2, and four slides from Grade 3 were selected for the analysis. An interstitial inflammation score of 0 means a pathologist estimated less than 10% of unscarred cortical parenchyma contained inflammation, while a score of 1 contains between 10-25%, a score of 2 contains between 26-50%, and a score of 3 contains >50%. Banff scores cover a variety of lesions, including but not limited to glomerulitis, tubulitis, interstitial fibrosis, tubular atrophy, and arteriolar hyalinosis. First, slides were segmented on an FTU-level, using our previously published and validated multicompartment segmentation algorithm^26^. Two new DigitAb models were trained for these external use cases. For both, the training set consisted of a combination of the IU-W frozen, IU-W FFPE, and IU-K datasets. The first model was trained only by supervision. Both the transplant and diabetic samples (described in next section) were then used as unlabeled samples for domain adaptation training for the second model. Performances were compared qualitatively as no ground truth labels were available. The transplant slides were then segmented using our domain-adaptive DigitAb model. By using the FTU segmentations, the cortical interstitial space was isolated from the rest of the biopsy, and several morphometric features were measured from the tissue. The percentage of tissue occupied by cortical interstitium was calculated, as well as both the count and area densities of immune cells in the cortical interstitium. The correlations between each of these values and the Banff inflammation score were represented by *R*^2^ values. Tubules segmented by the previously published FTU model were converted to specific segment definitions by assigning each a label from the most predicted cell type within the tubular boundary. Immune cells were matched to the closest FTU (glomeruli, arteries/arterioles, proximal tubules, thick ascending limbs, distal convoluted tubules, and collecting ducts) by KDTrees^50^ and placed in bins. To assess whether the distribution of immune cells in bins varied with inflammation score, analyses were conducted at the patient level, and inflammation score was treated as an ordered variable. Each patient was associated with a single outcome category reflecting increasing inflammation score. For each bin, a monotonic trend analysis was performed to test whether patient-level FTU proportions increased or decreased with inflammation score, measured by Kendall’s rank correlation coefficient^30^. This provides a non-parametric measure of association, and statistical significance was assessed using two-sided tests with alpha of 0.05.

### Use-Case 2: Immune Cell Distribution in Diabetic Nephropathy

Kidney biopsies from healthy control patients, and patients with diabetic nephropathy were obtained from the pathology archives of WUSTL Kidney Translational Research Center (KTRC). Patient metadata includes age, sex, race, estimated glomerular filtration rate (eGFR), and serum creatinine (SCr) during biopsy and follow-up for up to 3 years. A cohort of 135 slides were segmented twice, using one non-domain-adapted model, and one domain-adapted model. Since there were no cell type ground truths available for this cohort, segmentation results were assessed qualitatively by visual inspection. These samples were also segmented by the previously mentioned multicompartment segmentation algorithm, and immune cell densities (from the domain-adapted model output) were measured for this cohort as well. These values were correlated with eGFR, quantified by the coefficient of determination, *R^2^*. The same immune cell proximity testing was performed in this dataset, but instead of inflammation scores we use 3-year composite outcome, defined as either a doubling of serum creatinine, 50% decline in eGFR, or progression to end stage kidney disease within 3 years of biopsy. For this binary outcome, patient-level FTU bin proportions were compared between outcomes using non-parametric Mann-Whitney U tests for each bin. Additionally, a global test of compositional differences across all FTU bins was performed using a permutation-based multivariate approach. This global MANOVA assessed the difference in overall distribution across FTUs from patients with different outcomes. Statistical significance was assessed at an alpha of 0.05. We conducted a simulation-based power analysis of patient level FTU distribution to assess if limited sample size could explain the lack of statistical significance. A global test statistic was defined as the Euclidean distance between the mean bin proportion vectors between outcomes. Outcome labels were randomly assigned across patients to generate a null distribution. Power was then estimated using Monte Carlo simulation, assuming that the observed multivariate effect remains stable with increasing sample size. Patient-level proportion vectors were then resampled with replacement from the existing cohort, the global permutation test was applied, and the proportion of simulations with *p* < 0.05 were reported as the power.

## Acknowledgements

The authors acknowledge the University of Florida Research Computing for providing computational resources and support that have contributed to the research results reported in this publication. T.M.E., S.J., M.T.E. were supported by U54 DK134301. PS’s work was funded by NIH funding from NIDDK— R21DK128668, R01 DK114485, R01AR080668, R01DK129541, OT2OD038014, and from OD OT2 OD033753. The authors also thank Amanda Knoten for tissue processing.

## Author Contributions

P.S. and S.J. conceived the idea, S.J. T.M.E., P.S. and N.L. developed execution plan. Providing tissue samples, and Phenocycler and brightfield histology imaging were done by A.S., D.B., M.F., W.B., M.F., K.Y.J., M.T.E., A.S., K.C. Domain expertise was provided by S.J. and T.E.M. Analysis was done by N.L., S.W., T.M.E., S.J., P.S. Code was written by N.L. and S.W. Online Plugin was developed by A.T. N.L. wrote the manuscript, and P.S. mentored him during the process.

## Data and Code Availability

Access to images used in this study can be viewed at https://athena.rc.ufl.edu. Instructions and figures for navigating the data and running the segmentation plugin are outlined in **Supplementary Information**. Plugins for the digital slide archive are deployed as a docker image, which includes all codes and dependencies.^31^ All model files are made available for segmentation tasks. The codes used in this project can be found at https://github.com/SarderLab/DigitAb_Master.

One sample is shared for each of the three labeled datasets, including both prediction and ground truth annotations. All samples are shared for the UCD transplant cohort, and six samples are shared for the Washington University DN cohort. Three of these are from patients who progressed to a composite outcome within three years post biopsy, and the other three are from those who did not. Composite outcome defined as either reaching end stage renal disease, serum creatinine doubling, eGFR declining by 50% from time of biopsy. Both additional datasets include prediction annotations.

