## Supplemental Document for "DigitAb: Domain-Adaptive Cell Type Prediction Method from Light Microscopy Images"

**List of Figures and Tables:**

1. **Supplementary Figure 1. Phenocycler Expressions Overlaid on Histological Sections.**
2. **Supplementary Figure 2. Subcluster Expression for Select Markers in IU-W FFPE Cluster 6**.
3. **Supplementary Figure 3. Segmentation Performances.**
4. **Supplementary Figure 4. HistoQC Analysis.**
5. **Supplementary Table 1. Patient Information for Fresh Frozen Dataset.**
6. **Supplementary Table 2. Patient Information for FFPE Dataset.**
7. **Supplementary Table 3. Patient Information for KPMP Healthy Reference Tissue.**
8. **Supplementary Table 4. Patient Information for UC Davis Transplant Biopsies.**
9. **Supplementary Table 5. Patient Information for Washington University Diabetic Cohort.**
10. **Supplementary Table 6. Phenocycler Antibody Markers: Kidney.**
11. **Supplementary Table 7. Classification Justifications in IU-W Frozen Dataset.**
12. **Supplementary Table 8. Classification Justifications in IU-W FFPE Dataset.**
13. **Supplementary Table 9. Classification Justifications in IU-K samples.**
14. **Supplementary Table 10. Kidney Class Structure.**
15. **Navigating Whole Slide Images (WSIs), Molecular Images, and Annotations in the Digital Slide Archive (DSA)**
    a. **Supplementary Figure 5. Histology Navigation in the Digital Slide Archive (DSA).**
    b. **Supplementary Figure 6. Phenocycler Navigation in the Digital Slide Archive (DSA).**
    c. **Supplementary Figure 7. Viewing Annotations in Histological Sections.**
    d. **Supplementary Figure 8. Finding the Segmentation Algorithm on the DSA.**
    e. **Supplementary Figure 9. Segmentation Algorithm Parameters.**

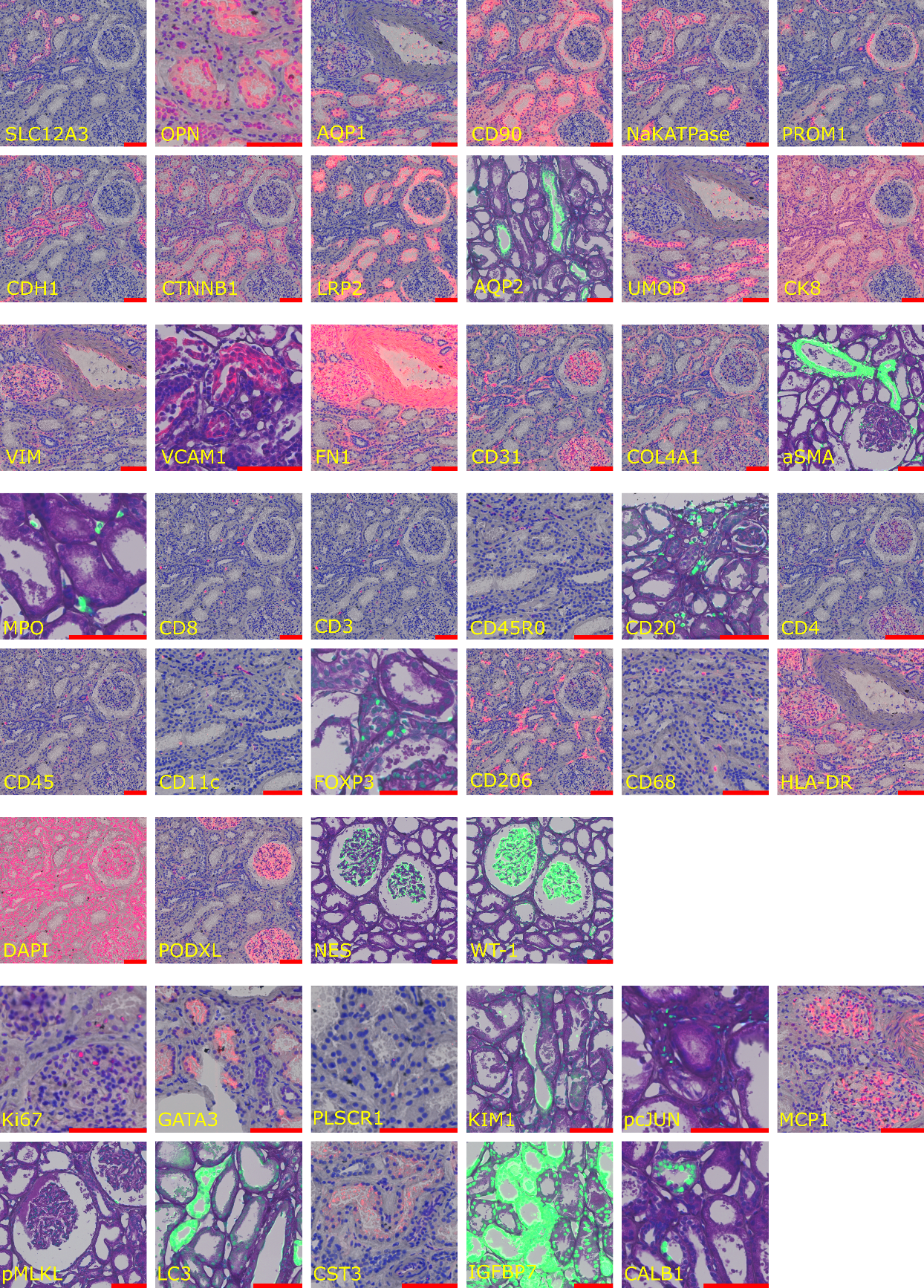

**Supplementary Figure 1. Phenocycler Expressions Overlaid on Histological Sections.**

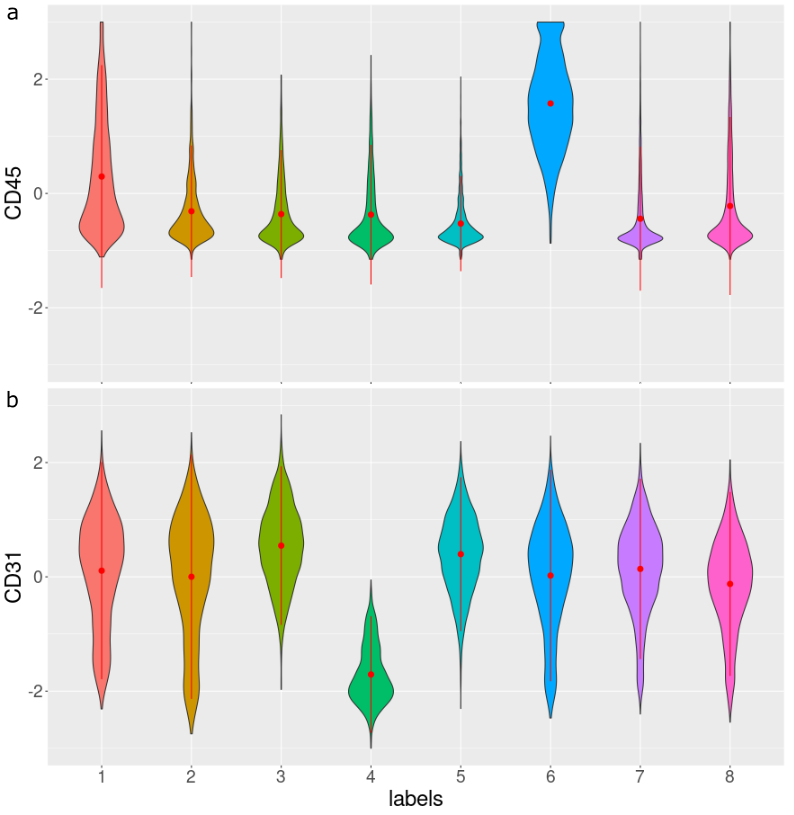

**Supplementary Figure 2. Subcluster Expression for Select Markers in IU-W FFPE Cluster 6**.

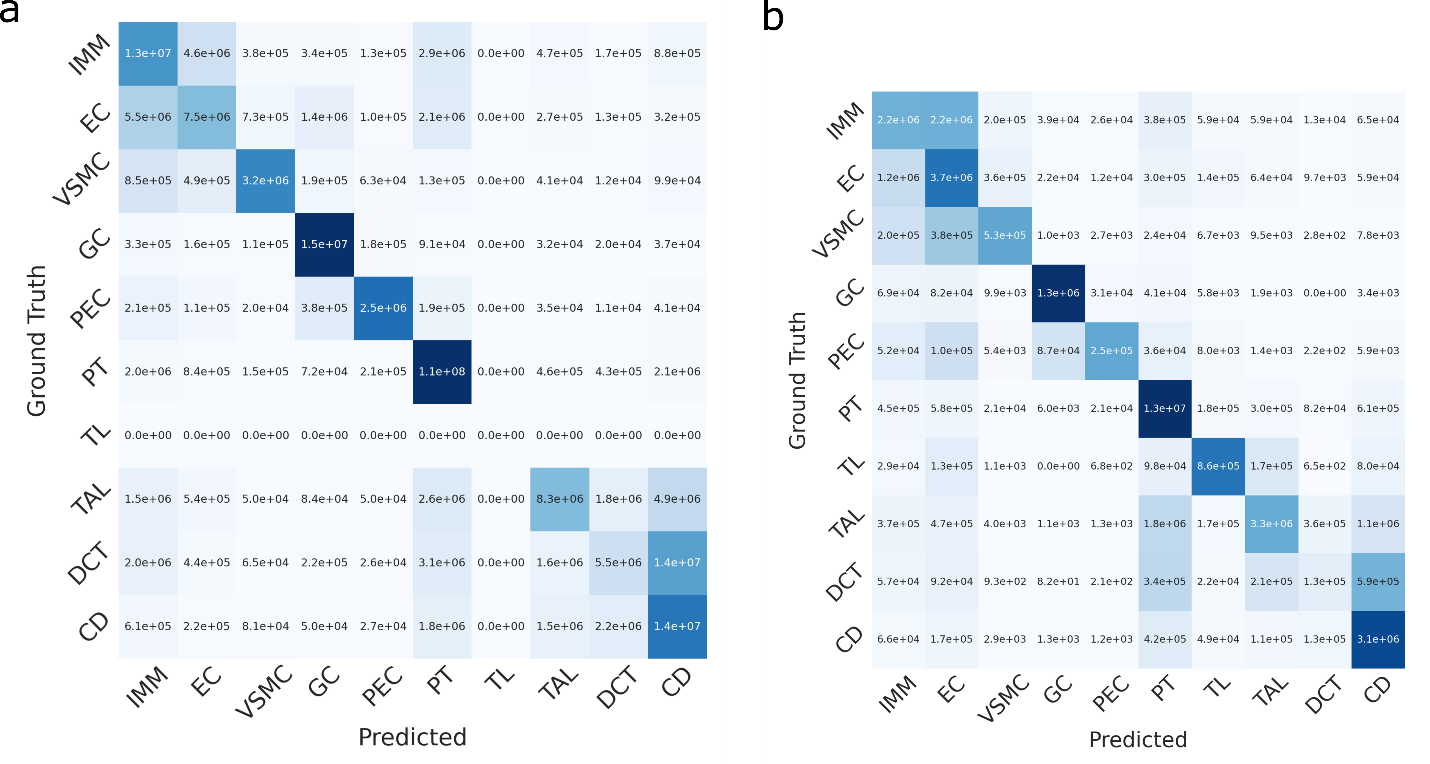

**Supplementary Figure 3. Segmentation Performances.** a) IU-W FFPE dataset. b) IU-K healthy reference tissue.

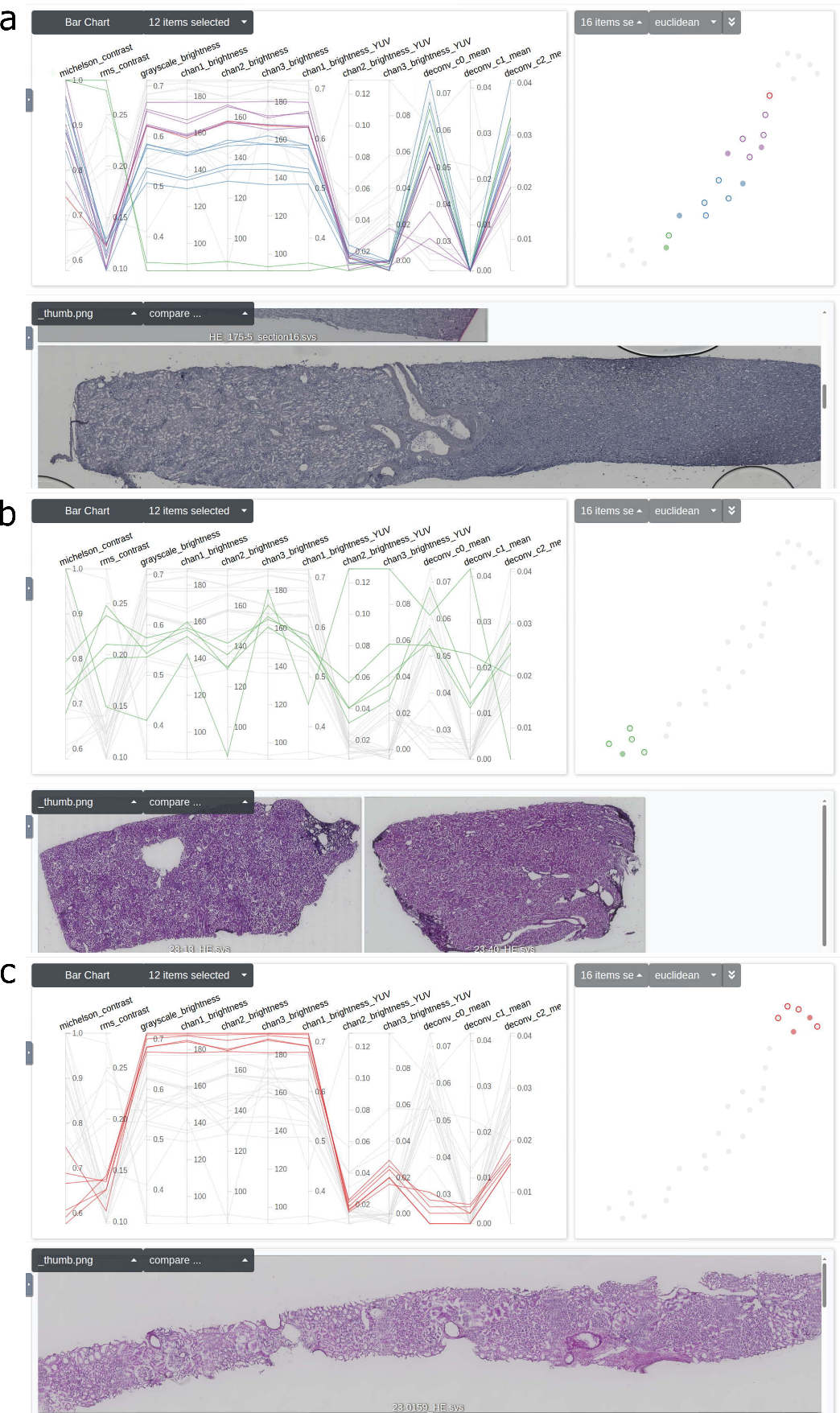

**Supplementary Figure 4. HistoQC Analysis.** a) IU-W frozen samples, b) IU-W FFPE samples, c) IU-K biopsies. Dimensionality reduction plots show distinct clustering of different cohorts.

**Supplementary Table 1. Patient Information for Fresh Frozen Dataset.**

| Slide Name | Participant ID | Tissue Source | Enrollment Category | Sex | Age (Years) | Race |
| --- | --- | --- | --- | --- | --- | --- |
| HE_5.svs | K2200007-3FB | HuBMAP | Healthy Reference | M | 46 | W |
| HE_6.svs | K2200007-3FB | HuBMAP | Healthy Reference | M | 46 | W |
| HE_7.svs | K2200007-3FB | HuBMAP | Healthy Reference | M | 46 | W |
| HE_175-5_section13.svs | K2200175-5PB | HuBMAP | Healthy Reference | M | 46 | W |
| HE_175-5_section15.svs | K2200175-5PB | HuBMAP | Healthy Reference | M | 46 | W |
| HE_175-5_section16.svs | K2200175-5PB | HuBMAP | Healthy Reference | M | 46 | W |
| HE_198_Section6.svs | K2100198_3 | HuBMAP | Healthy Reference | M | 51 | W |
| K22-256-5-6.svs | K2200256_5 | HuBMAP | Healthy Reference | F | 78 | W |
| K22-256-5-7.svs | K2200256_5 | HuBMAP | Healthy Reference | F | 78 | W |
| K22-256-5-8.svs | K2200256_5 | HuBMAP | Healthy Reference | F | 78 | W |
| K22-256-5-9.svs | K2200256_5 | HuBMAP | Healthy Reference | F | 78 | W |
| HE_0020_section1.svs | 21-0020 | HuBMAP | Healthy Reference | M | 70-79 | N/A |
| HE_0020_section2.svs | 21-0020 | HuBMAP | Healthy Reference | M | 70-79 | N/A |
| HE_0020_section3.svs | 21-0020 | HuBMAP | Healthy Reference | M | 70-79 | N/A |

**Supplementary Table 2. Patient Information for FFPE Dataset.**

| Participant ID | Tissue Source | Enrollment Category | Tissue Collection | Sex | Age | Race |
| --- | --- | --- | --- | --- | --- | --- |
| K21000184_7PB | HuBMAP | Healthy Reference | Deceased Donor | M | 60-69 | White |
| K2200013_4PB | HuBMAP | Healthy Reference | Deceased Donor | M | 40-49 | White |
| K2300040_12PB | HuBMAP | Healthy Reference | Deceased Donor | F | 67 |  |
| K2300080_6PB | HuBMAP | Healthy Reference | Deceased Donor | M | 38 |  |
| K2200038PB | HuBMAP | Healthy Reference | Nephrectomy | M | 40-49 | White |

**Supplementary Table 3. Patient Information for KPMP Healthy Reference Tissue.**

| Slide Name | Participant ID | Tissue Source | Enrollment Category | Sex | Age | Race |
| --- | --- | --- | --- | --- | --- | --- |
| 24-0168_HE.svs | 164-1 | KPMP | Healthy Reference | M | 50-59 | White |
| 24-0169_HE.svs | 164-5 | KPMP | Healthy Reference | F | 30-39 | White |
| 24-0170_HE.svs | 164-6 | KPMP | Healthy Reference | F | 30-39 | White |
| 23-0162_HE.svs | 164-12 | KPMP | Healthy Reference | F | 30-39 | White |
| 23-0159_HE.svs | 164-17 | KPMP | Healthy Reference | F | 30-39 | White |
| 23-0160_HE.svs | 164-21 | KPMP | Healthy Reference | F | 40-49 | White |
| 23-0158_HE.svs | 164-22 | KPMP | Healthy Reference | M | 20-29 | White |
| 24-0171_HE.svs | 165-13 | KPMP | Healthy Reference | F | 40-49 | Asian |

**Supplementary Table 4. Phenocycler Antibody Markers.**

| **Antibody** | **RRID** | **Target** | **Host** | **Vendor** | **Cat#** | **Clone** |
| --- | --- | --- | --- | --- | --- | --- |
| CD8 | AB_2895049 | CD8+ T cells | Mouse | Akoya Biosciences | 4150004 | SK1 |
| MPO | AB_2927678 | Granulocytic immune cells | Rabbit | Akoya Biosciences | 4250083 | E1E7I |
| CD3 | AB_2895047 | Pan T cells | Mouse | Akoya Biosciences | 4350008 | UCHT1 |
| CD20 | AB_2895056 | B cells | Mouse | Akoya Biosciences | 4150018 | L26 |
| CD45RO | AB_2895053 | Memory T cells | Mouse | Akoya Biosciences | 4250023 | UCHL1 |
| CD4 | AB_2895048 | CD4+ T cells | Mouse | Akoya Biosciences | 4350010 | SK3 |
| CD45 | AB_2895052 | Leukocytes | Mouse | Akoya Biosciences | 4150003 | HI30 |
| Ki67 | AB_2895046 | Proliferative cells | Mouse | Akoya Biosciences | 4250019 | B56 |
| CD11c | AB_2895050 | Dendritic cells | Mouse | Akoya Biosciences | 4350012 | S-HCL-3 |
| Vimentin | AB_393716 | Fibroblasts | Mouse | BD Pharmingen | 550513 | RV202 |
| GATA3 | AB_2927677 | T helper 2 cells | Rabbit | Akoya Biosciences | 4250085 | D13C9 |
| FOXP3 | AB_2927679 | Immune/T-reg cells | Mouse | Akoya Biosciences | 4550071 | 236A/E7 |
| OPN | AB_2194997 | Injury | Mouse | Santa Cruz Biotechnology | sc21742 | AKm2A1 |
| SLC12A3 | AB_571116 | Distal convoluted tubule epithelium | Rabbit | MiliporeSigma | AB3553 | Polyclonal |
| PLSCR1 | AB_2610379 | Injury | Mouse | Thermo Fisher Scientific | MA5-19636 | 3B4 |
| VCAM1 | AB_2895043 | Injury, Proximal tubule epithelium failed repair | Rabbit | Abcam | ab271899 | EPR5047 |
| CD206 | AB_2063019 | Immune, Mononuclear phagocytic cell | Goat | R&D Systems | AF2534 | Polyclonal |
| CD90 | AB_2895055 | Proximal tubule, Fibroblasts, Activated endothelial cells | Mouse | Akoya Biosciences | 4150021 | 5E10 |
| AQP1 | AB_2895040 | Proximal tubule epithelium | Mouse | Santa Cruz Biotechnology | sc-32737-X | 1/22 |
| KIM-1 | AB_2116559 | Injury | Mouse | R&D Systems | MAB1750 | 219211 |
| CCL2 | AB_626820 | Injury | Mouse | Santa Cruz Biotechnology | sc-32771 | 5J |
| p-c-Jun | AB_2895041 | Injury | Rabbit | Cell Signalling Technology | 3270BF | D47G9 |
| p-MLKL | AB_2895044 | Necroptosis | Rabbit | Cell Signaling Technology | 91689BF | D6H3V(S358) |
| Fibronectin | AB_2895045 | Injury, Fibroblasts | Rabbit | Abcam | ab271831 | F1 |
| LC3 | AB_1079382 | Autophagy | Rabbit | Sigma-Aldrich | L8918-25UL | Polyclonal |
| CST3 | AB_2895067 | Injury | Rabbit | Abcam | ab217569 | EPR4413 |
| ATP1A1 | AB_2888530 | Distal convoluted tubule epithelium | Rabbit | GeneTex | GTX635461 | HL114 |
| CD31 | AB_2895051 | Endothelial cells | Mouse | Akoya Biosciences | 4250009 | WM59 |
| PROM1 | AB_244339 | Injury | Mouse | Miltenyi Biotec | 130-090-422 | AC133 |
| Podocalyxin | AB_354920 | Podocytes | Goat | R&D Systems | AF1658 | Polyclonal |
| E-cadherin | AB_2895057 | Distal convoluted tubules, Collecting duct, loop of Henle | Mouse | Akoya Biosciences | 4250021 | 4A2C7 |
| IGFBP7 | AB_1617562 | Injury | Rabbit | OriGene | AP01109PU-S | Polyclonal |
| LRP2 | AB_2927680 | Proximal tubule epithelium | Mouse | R&D Systems | MAB9578 | 545606 |
| β-catenin | AB_2895058 | Tubule epithelium | Mouse | Akoya Biosciences | 4450036 | 12F7 |
| COL4A1 | AB_2927676 | Injury, Fibrosis | Rabbit | Akoya Biosciences | 4550122 | EPR209660 |
| AQP2 | AB_11154667 | Collecting duct principal cells | Rabbit | Thermo Fisher Scientific | PA5-22865 | Polyclonal |
| α-SMA | AB_2572996 | Vessels, Fibroblasts | Mouse | Thermo Fisher Scientific | 14-9760-82 | 1A4 |
| CD68 | AB_11151139 | Immune, Myeloidal, and Macrophage | Mouse | Thermo Fisher Scientific | 14-0688-82 | KP1 |
| Uromodulin | AB_2212386 | Thick ascending limb | Sheep | R&D Systems | AF5144 | Polyclonal |
| HLA-DR | AB_2895054 | Antigen presenting cells | Mouse | Akoya Biosciences | 4250006 | L243 |
| Cytokeratin-8 | AB_2895039 | Collecting duct epithelium | Mouse | Novus Biologicals | NBP2-34501 | TS1 |
| Nestin | AB_525739 | Podocytes | Mouse | NovusBio | NB300-266 | 10C2 |
| Calbindin | AB_1658451 | Distal convoluted tubule epithelium | Mouse | Abcam | ab82812 | CB-955 |
| CD123 | AB_314576 | Dendritic cells | Mouse | BioLegend | 306002 | 6H6 |
| WT-1 | AB_905863 | Podocytes | Mouse | NovusBio | NB110-60011 | 6F-H2 |
| TUBB3 | AB_2313773 | Proliferative cells | Mouse | BioLegend | 801201 | TUJ1 |
| CD56 | AB_2922611 | NK cells, CD56+ T cells | Mouse | BioLegend | 380702 | A19063A |

**Supplementary Table 5.** **Patient Information for UC Davis Transplant Biopsies.**

| Slide Name | Bx.Reason | g | cg | mm | ci | ct | i | ti | t | ptc | ah | aah | cv | v | Age | Sex | Race | Height | Weight | SCr |
| --- | --- | --- | --- | --- | --- | --- | --- | --- | --- | --- | --- | --- | --- | --- | --- | --- | --- | --- | --- | --- |
| 54225 | Baseline | 0 | 0 | 0 | 1 | 1 | 0 | 0 | 0 | 0 | 3 | 3 | 2 | 0 | 47 | M | White | 183 | 124 | 1.2 |
| 54228 | Baseline | 0 | 0 | 0 | 1 | 1 | 2 | 2 | 2 | 0 | 0 | 0 | 3 | 0 | 26 | F | Hispanic | 163 | 45.7 | 0.8 |
| 54240 | Baseline | 1 | 0 | 0 | 0 | 0 | 1 | 1 | 1 | 2 | 0 | 0 | 0 | 0 | 24 | F | White | 152 | 71 | 2.2 |
| 54246 | Baseline | 0 | 0 | 0 | 2 | 2 | 2 | 2 | 0 | 0 | 0 | 0 | 0 | 0 | 63 | M | Asian | 168 | 72.3 | 1.38 |
| 54397 | Baseline | 0 | 0 | 0 | 0 | 0 | 0 | 0 | 0 | 0 | 2 | 2 | 1 | 0 | 49 | M | White | 178 | 67.7 | 0.89 |
| 54403 | Baseline | 0 | 0 | 0 | 0 | 0 | 0 | 0 | 0 | 0 | 0 | 0 | 0 | 0 | 24 | M | Black | 157.48 | 69 | 2.98 |
| 54427 | Baseline | 1 | 0 | 0 | 0 | 0 | 2 | 2 | 1 | 1 | 1 | 1 | 2 | 0 | 21 | M | White | 175.26 | 84.5 | 0.8 |
| 54429 | Baseline | 2 | 0 | 0 | 0 | 0 | 1 | 1 | 0 | 2 | 0 | 0 | 0 | 0 | 14 | M | White | 172.72 | 67.9 | 1.2 |
| 61622 | Baseline | 0 | 0 | 0 | 0 | 0 | 0 | 0 | 0 | 0 | 2 | 2 | 1 | 0 | 49 | M | White | 178 | 67.7 | 0.89 |
| 61652 | Baseline | 2 | 0 | 0 | 0 | 0 | 1 | 1 | 0 | 2 | 0 | 0 | 0 | 0 | 14 | M | White | 172.72 | 67.9 | 1.2 |
| 62335 | Baseline | 0 | 0 | 0 | 0 | 0 | 1 | 1 | 0 | 1 | 0 | 0 | 0 | 0 | 0 | F | Black | 71 | 8.6 | 1.5 |
| 69733 | Baseline | 0 | 0 | 0 | 0 | 0 | 0 | 0 | 0 | 0 | 0 | 0 | 0 | 0 | 33 | F | Hispanic | 152.4 | 63 | 5.2 |
| 70141 | Baseline | 0 | 0 | 0 | 0 | 0 | 1 | 1 | 0 | 0 | 2 | 2 | 0 | 0 | 37 | M | White | 178 | 76.5 | 6.6 |
| 105692 | Cause | 2 | 0 | 0 | 0 | 0 | 3 | 3 | 2 | 0 | 1 | 2 | 2 | 0 | N/A | N/A | N/A | N/A | N/A | N/A |
| 105695 | Cause | 0 | 0 | 0 | 0 | 0 | 3 | 3 | 2 | 2 | 2 | 3 | 2 | 0 | N/A | N/A | N/A | N/A | N/A | N/A |
| 105704 | Cause | 0 | 0 | 0 | 0 | 0 | 3 | 3 | 3 | 0 | 0 | 0 | 0 | 0 | N/A | N/A | N/A | N/A | N/A | N/A |
| 105710 | Cause | 2 | 1a | 0 | 2 | 2 | 3 | 3 | 2 | 2 | 1 | 2 | 2 | 0 | N/A | N/A | N/A | N/A | N/A | N/A |
| 110904 | Time zero (implantation) | 0 | 0 | 0 | 0 | 0 | 2 | 2 | 1 | 0 | 1 | 2 | 0 | 0 | N/A | N/A | N/A | N/A | N/A | N/A |
| 112020 | Time zero (implantation) | 1 | 0 | 0 | 0 | 0 | 2 | 2 | 1 | 0 | 0 | 0 | 0 | 0 | 52 | F | White | 160 | 89.2 | 0.6 |

**Supplementary Table 6. Patient Information for Washington University Diabetic Cohort.**

| Specimen Label | Dx | Age | Sex | Race | eGFR | SCr | 3y Composite |
| --- | --- | --- | --- | --- | --- | --- | --- |
| K1300479 | DN | 26 | F | W | 34.5 | 2.04 | 1 |
| K1700183 | DN | 71 | M | W | 51.07 | 1.48 | 0 |
| K1300473 | DN | 41 | F | AA | 45.13 | 1.51 | 0 |
| K1300472 | DN | 63 | M | W | 12.72 | 7.11 | 1 |
| K1300471 | DN | 69 | M | W | 31.89 | 2.37 | 0 |
| K1300468 | DN | 61 | M | AA | 15.75 | 5.64 | 1 |
| K1300466 | DN | 54 | F | AA | 21.21 | 3.21 | 1 |
| K1600021 | HC | 56 | F | W | 42.38 | 1.5 | 0 |
| K1600289 | HC | 70 | M | AA | 56.58 | 1.41 | 0 |
| K1600292 | HC | 73 | M | W | 38.96 | 1.79 | 0 |
| K1300474 | DN | 34 | F | AA | 18.17 | 4.69 | 1 |
| K1700396 | DN | 69 | F | AA | 7.6 | 7.52 | 1 |
| K1800057 | DN | 15 | M | W | 52 | 1.87 | 0 |
| K1800059 | DN | 46 | F | AA | 35.86 | 2.81 | 0 |
| K1600370 | DN | 34 | F | W | 56.4 | 2.25 | 0 |
| K1600544 | HC | 63 | M | W | 57 | 1.26 | 0 |
| K1700251 | DN | 39 | M | AA | 21 | 3.35 | 0 |
| K1700383 | DN | 61 | F | AA | 14.96 | 4.81 | 0 |
| K1700524 | DN | 26 | F | AA | 20.2 | 4.37 | 1 |
| K1700528 | HC | 44 | M | W | 70.33 | 1.14 | 0 |
| K1800024 | HC | 62 | M | W | 59 | 1.26 | 0 |
| K1300477 | DN | 61 | M | W | 25.5 | 2.97 | 0 |
| K1900262 | DN | 41 | F | AA | 9.667 | 6.33 | 1 |
| K1900263 | DN | 52 | M | W | 20.5 | 3.45 | 0 |
| K1300470 | DN | 41 | F | AA | 31 | 3.29 | 0 |
| K1900264 | DN | 58 | M | AA | 49.11 | 3.07 | 0 |
| K1900265 | DN | 72 | M | W | 30 | 2.1 | 0 |
| K1800200 | DN | 33 | F | AA | 58.5 | 1.48 | 0 |
| K1800212 | DN | 64 | F | W | 59 | 1 | 0 |
| K1700021 | DN | 29 | M | AA | 51.14 | 1.93 | 0 |
| K1800022 | DN | 60 | F | AA | 41 | 1.67 | 0 |
| K1900215 | DN | 50 | F | A | 25.25 | 3.11 | 1 |
| K1900216 | DN | 29 | F | W | 23.2 | 3.08 | 0 |
| K1900218 | DN | 55 | M | W | 24.33 | 3.22 | 0 |
| K1900220 | DN | 58 | M | W | 18.6 | 3.84 | 1 |
| K1900221 | DN | 46 | F | AA | 24.75 | 2.69 | 0 |
| K1900223 | DN | 70 | M | W | 96 | 0.69 | 0 |
| K1900224 | DN | 61 | M | AA | 57.25 | 1.72 | 1 |
| K1900226 | DN | 69 | F | AA | 27 | 2.61 | 1 |
| K1900227 | DN | 48 | F | AA | 65.8 | 1.02 | 0 |
| K1900228 | DN | 42 | F | AA | 12.67 | 5.36 | 1 |
| K1900229 | DN | 49 | F | AA | 67.6 | 1.12 | 0 |
| K1900230 | DN | 58 | M | AA | 11.67 | 6.54 | 1 |
| K1900256 | DN | 69 | M | AA | 35.33 | 2.58 | 0 |
| K1900257 | DN | 61 | M | AA | 54.86 | 2.04 | 0 |
| K1900258 | DN | 68 | F | W | 24.17 | 2.2 | 0 |
| K1900259 | DN | 77 | M | W | 12 | 5.65 | 1 |
| K1900260 | DN | 71 | M | W | 36 | 2.03 | 0 |
| K1900351 | DN | 56 | F | AA | 21.95 | 3.34 | 1 |
| K1900441 | DN | 62 | M | W | 49.8 | 1.59 | 0 |
| K1900442 | DN | 54 | F | W | 29.15 | 2.09 | 0 |
| K2000151 | DN | 60 | M | W | 48.05 | 1.79 | 1 |
| K2000154 | DN | 70 | F | W | 18.15 | 3.91 | 1 |
| K2000145 | DN | 40 | F | W | 75.6 | 0.95 | 1 |
| K2000268 | DN | 75 | M | W | 19.85 | 3.04 | 0 |
| K2000286 | DN | 46 | F | AA | 14.35 | 3.85 | 1 |
| K2000292 | DN | 70 | F | W | 10.2 | 4.24 | 1 |
| K2000294 | DN | 72 | M | W | 10.25 | 6.08 | 1 |
| K2000295 | DN | 61 | M | AA | 22.65 | 3.73 | 0 |
| K2000298 | DN | 57 | M | W | 32.9 | 2.09 |  |
| K2000299 | DN | 72 | M | AA | 6.15 | 10.1 | 1 |
| K2000300 | DN | 72 | F | AA | 11.45 | 5.21 | 1 |
| K2000301 | DN | 42 | F | O/Pacific Islander/Hispanic | 53.4 | 1.29 | 0 |
| K2000307 | DN | 60 | M | O/Pacific Islander/Non-Hispanic | 23.75 | 2.98 | 0 |
| K2000256 | DN | 56 | M | AA | 141.4 | 0.7 | 0 |
| K2100024 | DN | 59 | F | AA | 44.05 | 1.6 | 0 |
| K2100025 | DN | 47 | F | AA | 68.75 | 1.06 | 0 |
| K1700444 | DN | 28 | M | W | 29.5 | 2.6 |  |
| K1800403 | DN | 25 | F | W | 41.4 | 1.53 |  |
| K1800429 | DN | 57 | F | W | 90.7 | 0.67 |  |
| K1900021 | DN | 45 | F | W | 62.8 | 0.96 |  |
| K1900069 | DN | 54 | M | W | 76.1 | 1.02 |  |
| K1900381 | DN | 66 | M | W | 96.7 | 0.8 |  |
| K2000089 | HC | 59 | F | W | 65 | 1 |  |
| K2000285 | DN | 52 | M | AA | 6.25 | 10.5 | 1 |
| K2000289 | DN | 80 | M | W | 28 | 2.2 |  |
| K2000302 | DN | 62 | F | AA | 24.57 | 2.24 | 0 |
| K2100049 | HC | 64 | F | W | 60 | 0.99 |  |
| K2100088 | DN | 63 | M | W | 91 | 1.06 | 0 |
| K2100138 | HC | 60 | M | W | 94 | 0.93 |  |
| K2100514 | DN | 50 | M | W | 30 | 2.42 |  |
| K2100518 | DN | 71 | M | W | 17 | 3.57 |  |
| K2100174 | HC | 65 | M | W | 97 | 0.84 |  |
| K2100197 | HC | 51 | M | W | 89 | 1.02 |  |
| K2100293 | DN | 63 | F | AA | 29.35 | 2.5 | 1 |
| K2100272 | HC | 53 | F | W | 69 | 1 |  |
| K2100287 | DN | 48 | M | AA | 22.25 | 3.75 | 1 |
| K2200027 | DN | 66 | F | AA | 21.67 | 2.65 | 1 |
| K2100529 | HC | 48 | M | W | 86 | 1.07 |  |
| K2200005 | HC | 46 | M | W | 102 | 0.9 |  |
| K2200084 | HC | 59 | F | W | 87 | 0.78 |  |
| K2200215 | HC | 46 | M | W | 91 | 1.03 |  |
| K2200189 | DN | 58 | M | AA | 7 | 11.9 | 1 |
| K2200190 | DN | 47 | F | W | 10 | 7.85 | 1 |
| K2200192 | DN | 70 | M | AA | 12.8 | 5.07 | 1 |
| K2200193 | DN | 60 | M | AA | 82.5 | 1.14 | 0 |
| K2200194 | DN | 56 | M | W | 22.5 | 3.07 | 1 |
| K2200195 | DN | 75 | F | AA | 22.29 | 2.23 | 0 |
| K2200196 | DN | 88 | F | AA | 87 | 0.57 |  |
| K2200197 | DN | 58 | M | AA | 26 | 2.72 |  |
| K2200198 | DN | 78 | M | W | 29 | 2.72 | 1 |
| K2200199 | DN | 52 | M | W | 14.94 | 4.68 | 1 |
| K2200200 | DN | 64 | F | W | 22.33 | 2.43 | 0 |
| K2200201 | DN | 64 | M | W | 25 | 2.63 | 0 |
| K2200204 | DN | 58 | M | A | 12 | 5.17 |  |
| K2200206 | DN | 53 | M | AA | 90 | 1 |  |
| K2200208 | DN | 43 | F | AA | 8.667 | 5.82 | 1 |
| K2200210 | DN | 58 | M | W | 25.33 | 3.15 |  |
| K2200211 | DN | 36 | F | AA | 34.08 | 2.22 | 1 |
| K2200212 | DN | 67 | M | AA | 11 | 5.6 | 1 |
| K2200240 | HC | 54 | M | AA | 100 | 0.91 |  |
| K2200253 | HC | 78 | F | W | 84 | 0.73 |  |
| K1900174 | HC | 76 | M | W | 55.1 |  |  |
| K1900317 | HC | 67 | M | W |  |  |  |
| K2100035 | HC | 65 | M | W | 48.75 | 1.58 |  |
| K2100102 | HC | 50 | M | W |  |  |  |
| K2100218 | HC | 57 | F | W |  |  |  |
| K2200414 | HC | 90 | M | W | 64 | 1.1 |  |
| K2300278 | HC | 69 | M | W | 57 | 1.2 |  |
| K2200539 | HC | 66 | M | AA | 65.5 | 1.33 |  |
| K2200573 | HC | 54 | M | W | 42 | 1.88 |  |
| K2300031 | HC | 61 | M | W | 52.33 | 1.55 |  |
| K2300023 | HC | 53 | F | W | 54.67 | 1.19 |  |
| K2300071 | HC | 30 | M | W | 57 | 1.65 |  |
| K2300241 | HC | 63 | F | W | 34 | 1.66 |  |
| K2300244 | HC | 49 | F | W | 58 | 1.17 |  |
| K2300287 | HC | 69 | F | W | 46.33 | 1.26 |  |
| K2300288 | HC | 22 | F | AA |  | 0.62 |  |
| K2300289 | HC | 45 | M | W |  |  |  |
| K2300290 | DN | 79 | F | AA | 30.5 | 1.72 |  |
| K2300291 | DN | 75 | F | AA | 7.25 | 6.06 |  |
| K2300292 | DN | 72 | F | W | 36.83 | 2.35 |  |
| K2300293 | DN | 63 | F | W | 72 | 0.9 |  |
| K2300294 | DN | 61 | F | AA | 30.75 | 1.95 |  |
| K2300295 | DN | 56 | F | W | 25 | 2.77 |  |
| K2300296 | DN | 49 | F | AA | 48.33 | 1.37 |  |
| K2300298 | DN | 40 | M | AA | 18.43 | 7.46 | 1 |
| K2300300 | DN | 40 | M | W | 52.5 | 1.72 |  |
| K2300302 | DN | 35 | M | AA | 30 | 2.77 |  |
| K1401965 | HC | 22 | M | AA | 59.33 | 1.46 |  |
| K2000291 | HC | 16 | M | AA |  |  |  |
| K2200383 | HC | 70 | F | W | 50.33 | 1.2 |  |
| K2300335 | HC | 20 | M | W |  |  |  |
| K2300336 | HC | 24 | M | W |  |  |  |
| K2300337 | HC | 29 | F | AA | 118 | 0.7 |  |
| K2300623 | HC | 6 | F | W |  | 0.33 |  |
| K2300624 | HC | 12 | M | W |  | 0.51 |  |

**Supplementary Table 7. Classification Justifications in IU-W Frozen Dataset.**

| Cluster | Classification | Upregulated Markers | Spatial Localization |
| --- | --- | --- | --- |
| 1 | Artifact | N/A | Edge of CODEX section |
| 2 | Artifact | N/A | Edge of CODEX section |
| 3 | Medullary immune + endothelial | CD45, CD31, CD206 | Medullary interstitium |
| 4 | Medullary endothelial | CD31 | Medullary interstitium |
| 5 | Cortical immune | CD45, CD206, CD68 | Cortical interstitium |
| 6 | Artifact | N/A | Vessel lumen debris |
| 7 | Proximal tubule epithelium | CK8, LRP2 | Tubular epithelium |
| 8 | Thin limb epithelium | CK8, distance to glom | Medullary epithelium |
| 9 | Medullary endothelial | CD31, FIB | Medullary interstitium |
| 10 | Thin limb epithelium | CK8 | Medullary epithelium |
| 11 | Medullary thick ascending limb epithelium | CK8, UMOD, NaK | Medullary epithelium |
| 12 | Medullary collecting duct epithelium | CK8, ECAD | Medullary epithelium |
| 13 | Medullary thick ascending limb epithelium | CK8, UMOD, NaK | Medullary epithelium |
| 14 | Thin limb epithelium | CK8 | Medullary epithelium |
| 15 | Cortical thick ascending limb epithelium | CK8, UMOD, NaK | Cortical epithelium |
| 16 | Proximal tubule epithelium | CK8, LRP2, AQP1, CD90 | Cortical epithelium |
| 17 | Proximal tubule epithelium | CK8, LRP2, AQP1 | Cortical epithelium |
| 18 | Cortical collecting duct epithelium | CK8, ECAD | Cortical epithelium |
| 19 | Vascular smooth muscle cell | FIB | Tunica media |
| 20 | Cortical immune | CD45, CD3, CD4, CD8 | Cortical interstitium |
| 21 | Vascular smooth muscle cell | FIB, VIM | Tunica media |
| 22 | Cortical immune | CD45, HLA-DR, CD206 | Cortical interstitium |
| 23 | Outer medulla collecting duct epithelium | CK8, ECAD, BCAT, AQP2 | Medullary epithelium |
| 24 | Inner medulla collecting duct epithelium | CK8, ECAD, BCAT, AQP2 | Medullary epithelium |
| 25 | Thin limb epithelium | CK8, AQP1 | Medullary epithelium |
| 26 | Medullary immune | CD45, CD45R0, CD206 | Medullary interstitium |
| 27 | Glomerular + parietal epithelium | CD31, PODXL, FIB, VIM | Glomerulus |
| 28 | Distal convoluted tubule epithelium | CK8, SLC12A3 | Cortical epithelium |
| 29 | Cortical distal convoluted tubule + thick ascending limb | CK8, SLC12A3, UMOD | Cortical epithelium |
| 30 | Proximal tubule epithelium | CK8, LRP2 | Cortical epithelium |
| 31 | Glomerular + parietal epithelium | CD31, PODXL, FIB, VIM | Glomerulus |

| Cluster 3 Subclusters | Classification | | Upregulated Markers | |
| --- | --- | --- | --- | --- |
| 1 | Endothelial | | CD31 | |
| 2 | Endothelial | | CD31 | |
| 3 | Immune | | CD45 | |
| 4 | Immune | | CD45 | |
| 5 | Endothelial | | CD31 | |
| 6 | Immune | | CD45 | |
| 7 | Immune | | CD45 | |
| 8 | Immune | | CD45 | |
| Cluster 27 Subclusters | | Classification | | Upregulated Markers |
| 1 | | Parietal epithelium | | PROM1 |
| 2 | | Glomerular | | PODXL |
| 3 | | Glomerular | | PODXL |
| 4 | | Glomerular | | PODXL |
| 5 | | Glomerular | | PODXL |
| 6 | | Parietal epithelium | | PROM1 |
| 7 | | Glomerular | | PODXL |
| 8 | | Glomerular | | PODXL |
| 9 | | Glomerular | | PODXL |
| 10 | | Glomerular | | PODXL |
| 11 | | Glomerular | | PODXL |

**Supplementary Table 8. Classification Justifications in IU-W FFPE Dataset.**

| Cluster | Classification | Upregulated Markers | Spatial Localization |
| --- | --- | --- | --- |
| 1 | Proximal tubule epithelium | CK8, LRP2 | Tubular epithelium |
| 2 | Proximal tubule epithelium | CK8, LRP2 | Tubular epithelium |
| 3 | Collecting duct epithelium | CK8, AQP2 | Tubular epithelium |
| 4 | Vascular smooth muscle cell | aSMA, VIM | Tunica media |
| 5 | Endothelium | CD31 | Interstitium |
| 6 | Endothelium + immune | CD31, CD45 | Interstitium |
| 7 | Artifact | N/A | Tissue edges and folds |
| 8 | Parietal epithelium | VCAM1, PROM1 | Glomerulus |
| 9 | Immune | CD45, CD45R0, CD206 | Interstitium |
| 10 | Thick ascending limb epithelium | CK8, UMOD | Tubular epithelium |
| 11 | Glomerular tuft | PODXL, CD31, WT1, NEST | Glomerulus |
| 12 | Proximal tubule epithelium | CK8, LRP2 | Tubular epithelium |
| 13 | Proximal tubule epithelium | CK8, LRP2 | Tubular epithelium |
| 14 | Proximal tubule epithelium | CK8, LRP2 | Tubular epithelium |
| 15 | Artifact | N/A | Lumen debris |
| 16 | Proximal tubule epithelium | CK8, LRP2 | Tubular epithelium |
| 17 | Distal convoluted tubule epithelium | CK8, SLC12A3 | Tubular epithelium |
| 18 | Proximal tubule epithelium | CK8, LRP2 | Tubular epithelium |
| 19 | Thick ascending limb epithelium | CK8, UMOD | Tubular epithelium |

| Cluster 6 Subclusters | Classification | Upregulated Markers |
| --- | --- | --- |
| 1 | Immune | CD45 |
| 2 | Endothelial | CD31 |
| 3 | Endothelial | CD31 |
| 4 | Enothelial | CD31 |
| 5 | Endothelial | CD31 |
| 6 | Immune | CD45 |
| 7 | Endothelial | CD31 |
| 8 | Endothelial | CD31 |

**Supplementary Table 9. Classification Justifications in IU-K samples.**

| Cluster | Classification | Upregulated Markers | Spatial Localization |
| --- | --- | --- | --- |
| 1 | Proximal tubule epithelium | CK8, LRP2, AQP1 | Tubular epithelium |
| 2 | Proximal tubule epithelium | CK8, LRP2, AQP1 | Tubular epithelium |
| 3 | Endothelial | CD31 | Tunica intima |
| 4 | Immune | CD45, CD11c, CD4, CD206 | Interstitium |
| 5 | Immune | CD45, CD11c, CD3, CD4, CD8 | Interstitium |
| 6 | Thick ascending limb epithelium | CK8, UMOD, NaK | Tubular epithelium |
| 7 | Endothelial | CD31, Nestin, WT1 | Tunica intima |
| 8 | Glomerular tuft | PODXL, CD31, Nestin, WT1 | Glomerulus |
| 9 | Thick ascending limb epithelium | CK8, UMOD, NaK | Tubular epithelium |
| 10 | Collecting duct epithelium | CK8, AQP2, ECAD | Tubular epithelium |
| 11 | Distal convoluted tubule epithelium | CK8, SLC12A3, NaK, ECAD | Tubular epithelium |
| 12 | Proximal tubule epithelium | CK8, LRP2, AQP1, PROM1 | Tubular epithelium |
| 13 | Vascular smooth muscle cell | aSMA | Tunica media |
| 14 | Thin limb | CK8, AQP1, PROM1 | Tubular epithelium |
| 15 | Parietal epithelium | PROM1, VCAM1, WT1 | Glomerulus |

**Supplementary Table 10. Kidney Class Structure.**

| Abbreviation | Cell Type |
| --- | --- |
| IMM | Immune Cell |
| EC | Endothelial Cell |
| SMC | Smooth Muscle Cell |
| GC | Glomerular Cell (Podocyte, EC-GC, MC) |
| PEC | Parietal Epithelial Cell |
| PT | Proximal Tubule Epithelial Cell |
| TL | Thin Limb Epithelial Cell (DTL, ATL) |
| TAL | Thick Ascending Limb Epithelial Cell |
| DCT | Distal Convoluted Tubule Epithelial Cell |
| CD | Collecting Duct Epithelial Cell |

**Navigating Whole Slide Images (WSIs), Molecular Images, and Annotations in the Digital Slide Archive (DSA)**

A User can log-in to the Digital Slide Archive (DSA; <https://athena.rc.ufl.edu/>) as a public user with the following credentials: *Username:* reviewer1; *password:* DigitAb2026!. The WSIs are located under *Collections/HuBMAP CODEX2Fusion/*.  This will bring the user to a page displaying each of the organs available for viewing. Upon clicking on an organ, the user may be presented with several cohorts of images, or in the case of only 1 available cohort, a choice between histology or molecular images. The molecular images contain unregistered Phenocycler images and will correspond to sections in the histology folder.

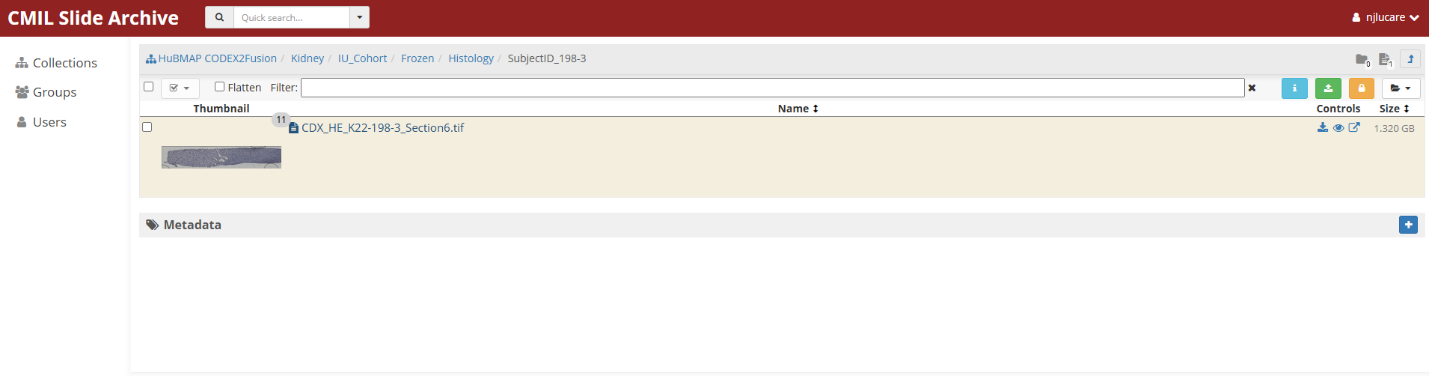

**Supplementary Figure 5. Histology Navigation in the Digital Slide Archive (DSA).** Within each dataset used in this manuscript, histology images are located under the *Histology* tab within *HuBMAP CODEX2Fusion*.

After navigating to the molecular data directory, and upon selecting an image and clicking on the file name, the user can see the image in a separate page. By clicking on the drop-down menu for the *Image control mode* and selecting *Channel Compositing*, the user will be able to see each of the marker layers in the Phenocycler image. Each of these layers can be toggled on and off, the display colors can be changed, and the range of values can be changed to increase or decrease the contrast. After clicking *Open in HistomicsUI* the user can zoom in and out and hover over different regions in the slide.  The user can again select *Channel Compositing* in the dropdown menu below *Frame Selector*.

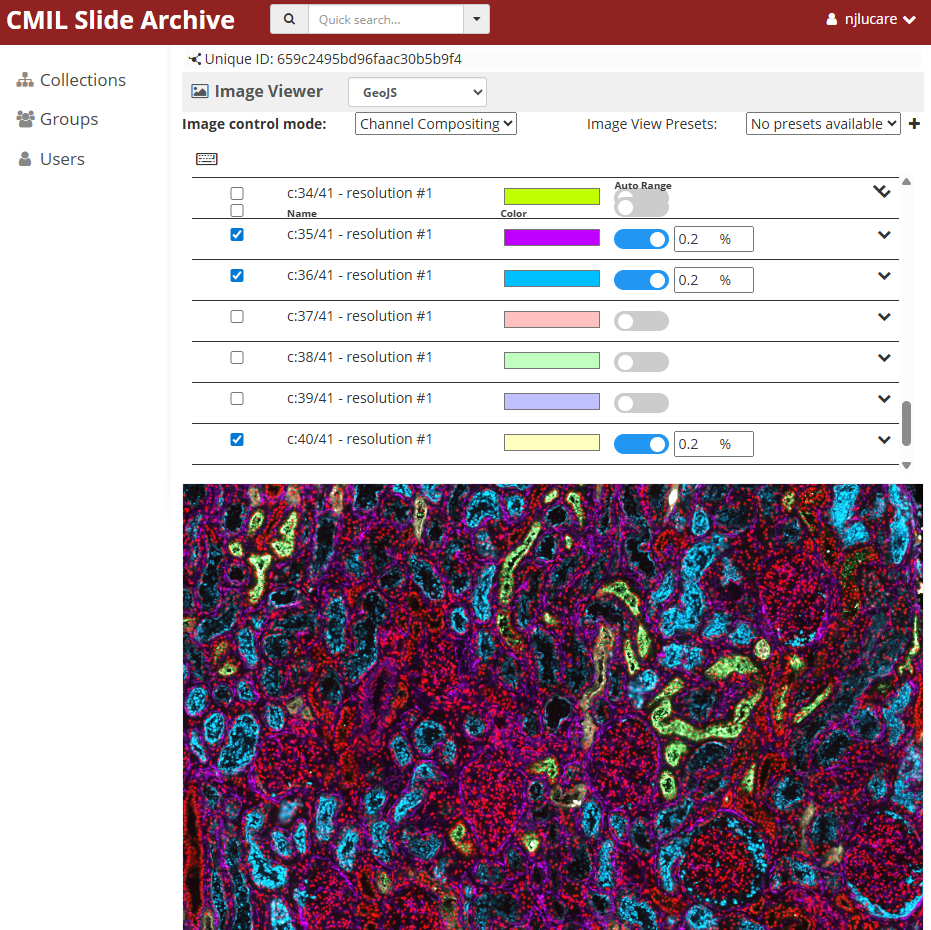

**Supplementary Figure 6. Phenocycler Navigation in the Digital Slide Archive (DSA).** Within each dataset used in this manuscript, Phenocycler spatial proteomic images are located under the *Molecular_Data* tab within *HuBMAP CODEX2Fusion*.

After navigating to the histology directory, and upon selecting a WSI and clicking on the file name, the user can see the WSI in a separate page with computational annotations in Json format and metadata associated with the image under the *Metadata* and *Annotations* section, respectively. After clicking *Open in HistomicsUI* the user can zoom in and out and hover over different regions in the slide. After clicking on the *Other* folder on the right panel, the ground truth annotations for each cell type become available, as well as artifacts that were discarded during performance analysis. These layers can be toggled on and off by selecting or deselecting the eye button. If many annotations need to be loaded, the contours may take some time to appear and may initially appear as circles in low-resolution viewing.

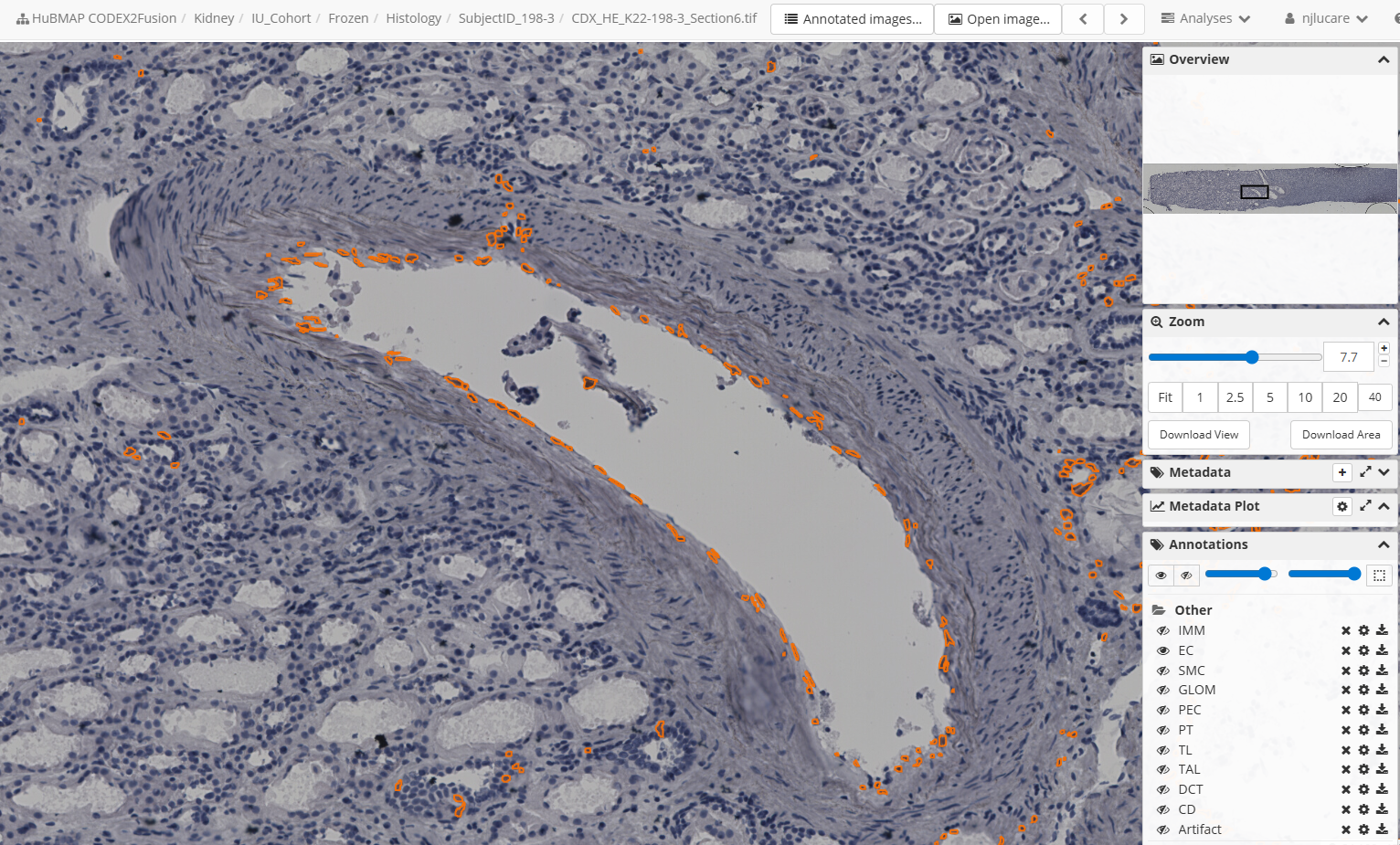

**Supplementary Figure 7. Viewing Annotations in Histological Sections.** By clicking on the *eye* icons next to each annotation layer, the contours can be toggled on and off for viewing.

Slides can be uploaded to user folders by selecting your username in the top right corner, clicking *My folders* then either creating or clicking on a folder. After clicking the green button in the top right, the user can upload their own slide. The segmentation algorithm can be run on the DSA platform by selecting the *Analyses* dropdown menu at the top right of the HistomicsUI interface. Navigate to *sarderlab/miscellaneous/Digitab Segmentation* > *Segment* to bring up the segmentation interface on the left side of the screen.

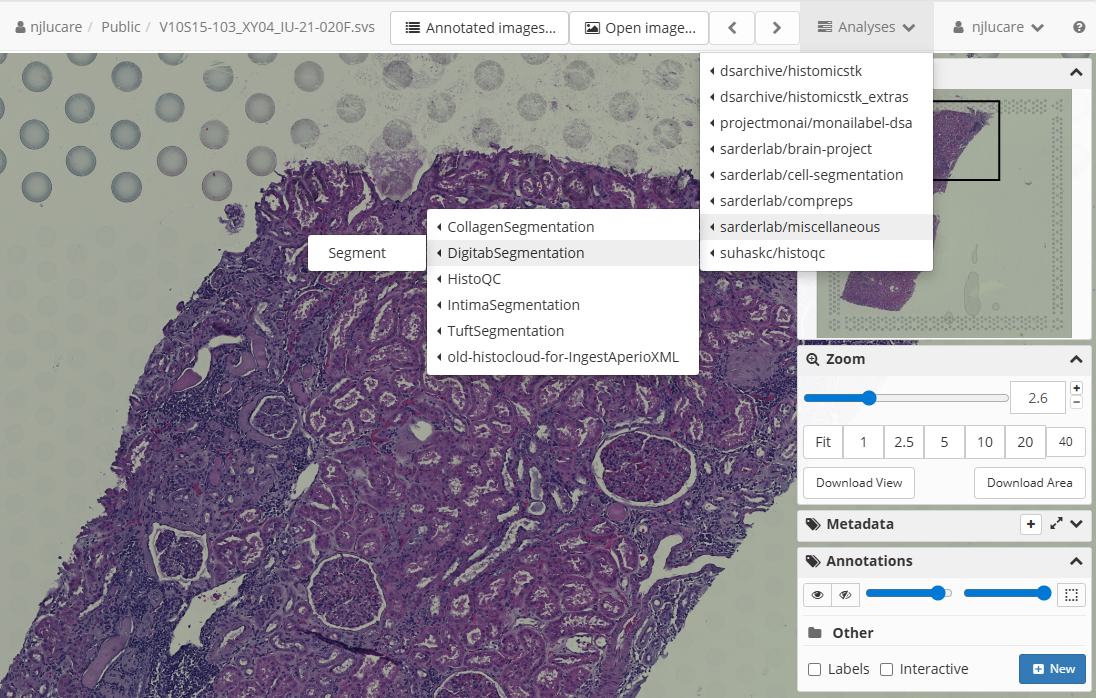

**Supplementary Figure 8. Finding the Segmentation Algorithm on the DSA.**

The opened image should already be selected, but you may select another image if desired. Choose the model file by selecting the folder icon and navigating to *Collections/HuBMAP CODEX2Fusion/model_files* and selecting a model file. Open the *Prediction Parameters* menu and set *Number of Classes* to 11 (10 cell types plus 1 background class). The other parameters may be adjusted to best segment your slide or optimized for the computational resources available.  For example, adjusting the confidence threshold will change the sensitivity of the network to background vs cell classification. For predictions on IU-W frozen samples, use the *ckpt_Frozen.pth* file in the *model_files* folder, with confidence threshold = 0.5, region size = 769, and step size = 512. Batch size may be changed but this will not affect the predictions. For predictions on IU-W FFPE samples, use the *ckpt_FFPE.pth* file in the same folder, and use the same parameters. For predictions on IU-K samples, use the *ckpt_KPMP.pth* file in the same folder, and use the same parameters.

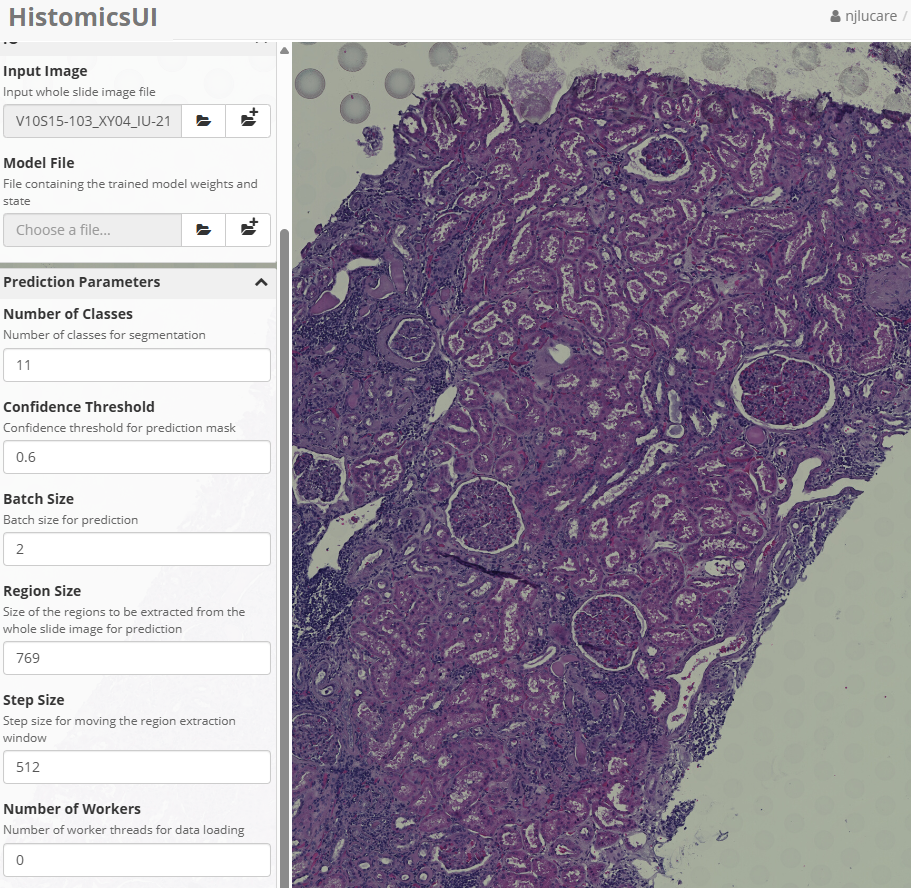

**Supplementary Figure 9. Segmentation Algorithm Parameters.** Number of classes should remain 11 for the available model files. Confidence threshold may be adjusted for over/under-segmentation in particular slides. Batch size will control time to completion, while region and step sizes control the ROI and stitching of patches.
